# SARS-CoV-2 nucleocapsid induces hyperinflammation and vascular leakage through the Toll-like receptor signaling axis in macrophages

**DOI:** 10.1101/2025.08.28.672752

**Authors:** Zhenlan Yao, Pablo A. Alvarez, Carolina Chavez, Yennifer Delgado, Prashant Kaushal, David Austin, Qian Li, Yanying Yu, Anne K Zaiss, Vaithilingaraja Arumugaswami, Qiang Ding, Jeffrey J. Hsu, Robert Damoiseaux, Mehdi Bouhaddou, Alexander Hoffmann, Melody M. H. Li

**Affiliations:** Department of Microbiology, Immunology and Molecular Genetics, University of California, Los Angeles, CA 90095, USA; Molecular Biology Institute, University of California, Los Angeles, CA 90095, USA; Department of Molecular and Medical Pharmacology, David Geffen School of Medicine at UCLA, Los Angeles, CA 90095, USA; Department of Medicine, University of California, Los Angeles, CA 90095, USA; Department for Bioengineering, Samueli School of Engineering, University of California, Los Angeles, CA 90095, USA; Department of Medicine, Veterans Affairs Greater Los Angeles Health Care System, Los Angeles, CA 90073, USA; California Nano Systems Institute, University of California, Los Angeles, CA 90095, USA; Institute for Quantitative and Computational Biosciences, University of California, Los Angeles, CA 90095, USA; Eli and Edythe Broad Center of Regenerative Medicine and Stem Cell Research, University of California, Los Angeles, CA 90095, USA; Center for Infection Biology, School of Basic Medical Sciences, Tsinghua University, Beijing, China

**Author notes:** Mass Spectrometry Technology Access Center at McDonnell Genome Institute, Washington University School of Medicine, St. Louis, MO, 63108, USA. These authors contributed equally (Zhenlan Yao, Pablo A. Alvarez).

## Abstract

Tens of thousands of severe COVID-19 cases are hospitalized weekly in the U.S., often driven by an imbalance between antiviral responses and inflammatory signaling, leading to uncontrolled cytokine secretion. The SARS-CoV-2 nucleocapsid (N) protein is a known immune antagonist, but its role in macrophage-driven cytokine storms is unclear. We demonstrate that N functions in a pathway-specific manner, specifically amplifying nuclear factor κB-related transcripts upon Toll-like receptor 7/8 stimulation. Moreover, we show that this is a conserved feature of pathogenic coronaviruses, with the delta variant N being the most pro-inflammatory. Our interaction networks suggest the delta variant N drives inflammation through interactions with several stress granule-related proteins. Profiling of secreted cytokines revealed that supernatants from the delta variant N-expressing macrophages disrupt brain and heart endothelial barriers, implicating N in COVID-19-associated cognitive and cardiac complications. Our findings highlight N-mediated immune imbalance as a driver of severe COVID-19 and identify N as a promising therapeutic target to mitigate hyperinflammation.

## INTRODUCTION

The emergence of severe acute respiratory syndrome coronavirus 2 (SARS-CoV-2) has caused significant global morbidity and mortality^1^. Even with the deployment of safe and effective vaccines, thousands of patients are hospitalized worldwide each winter^2^. Approximately one-third of these patients will develop acute respiratory distress syndrome (ARDS), for which there are currently no effective treatments^3,4^. It is well understood that ARDS is marked by unconstrained pro-inflammatory cytokine secretion^5^, termed cytokine storm. An additional hallmark of severe disease in these patients is a dampened antiviral response mediated by type I interferons (IFNs)^6,7^. Moreover, post-infectious sequelae often result in neurological and cardiac complications mediated by a dysregulated immune response. Together, these complications emphasize the need to elucidate mechanisms of SARS-CoV-2 immune dysregulation, yet our current understanding of these mechanisms remain controversial.

Type I IFNs, including IFN-α/β, are critical first line of defense against viral infections, including SARS-CoV-2. Their production is triggered by pattern recognition receptors (PRRs) that sense viral components within infected cells. In the case of SARS-CoV-2, retinoic acid-inducible gene I (RIG-I)-like receptors (RLRs) are considered the primary sensors of viral RNA^8^, though other PRRs, such as Toll-like receptors (TLRs), also contribute. Endosomal TLRs are highly expressed in immune cells, especially phagocytes, like monocytes, macrophages, and dendritic cells, and are responsible for sensing foreign viral RNA^9–11^. Importantly, these TLRs have been implicated in amplifying inflammation during SARS-CoV-2 infection^11,12^. Upon PRR activation, key transcription factors—including IFN regulatory factors 3 and 7 (IRF3/7) and nuclear factor κB (NF-κB)—translocate to the nucleus to induce antiviral and inflammatory gene expression. Secreted type I IFNs also engage the STAT1/STAT2/IRF9 complex to drive the expression of a broad array of IFN-stimulated genes (ISGs), many of which have direct antiviral functions. Timely induction of ISGs is essential for containing viral spread while delayed and impaired IFN responses are associated with severe COVID-19. While IFN signaling primarily drives antiviral defenses, NF-κB signaling promotes inflammation, and its sustained activation can lead to immunopathology. Indeed, dysregulated IFN and NF-κB responses are hallmarks of severe SARS-CoV-2 infection, underscoring the need to understand how the virus disrupts these pathways to evade immune control and promote disease.

Pathogenic coronaviruses, including Middle East respiratory syndrome coronavirus (MERS-CoV), SARS-CoV-1, and SARS-CoV-2 have several proteins known to interact with and dampen innate immune pathways^13–17^. Among them, the nucleocapsid (N) protein is consistently the most abundant immune modulator. Several N molecules are packaged within the virion and it is the most highly expressed protein during replication, with estimates reaching up to 10^8^ molecules per infected cell^18,19^. Moreover, N is reportedly found throughout the cell, including the nucleus, in addition to being presented on the cell surface^20,21^. Therefore, N proteins have several opportunities to interact with various cellular signaling components in infected and nearby uninfected cells. N is reported to associate with G3BP1/2 to inhibit the formation of stress granules, which are critical for promoting an antiviral state^22^. N has also been shown to impede the signaling of RLRs by blocking RIG-I interactions with critical cofactors to prevent downstream signaling^8,23^. Interestingly, newer SARS-CoV-2 variants have evolved mutations in N that better facilitate the manipulation of RLR signaling^24^, emphasizing the need to monitor the effects of N evolution on immune signaling.

As key players in innate immunity and crucial antigen-presenting cells for activating the adaptive immune system, macrophages are well-positioned to encounter SARS-CoV-2 N proteins, which are abundantly displayed on the surface of infected cells. Strikingly, macrophages are the predominant cell type responsible for the cytokine storm observed in severe COVID-19^25–27^. Specifically, monocyte-derived macrophages comprise a majority of the immune cells found in bronchoalveolar lavage fluid, further emphasizing their inflammatory role in SARS-CoV-2 infection^28^; however, direct infection of macrophages by SARS-CoV-2 remains controversial. Earlier studies suggest that macrophages are refractory to SARS-CoV-2 infection^29^; however, accumulating evidence so far reveals that SARS-CoV-2 infection depends on the macrophage subtype^26,30^. For example, interstitial macrophages allow productive SARS-CoV-2 replication, characterized by an altered transcriptome and dense viral RNA bodies^28,30^. While SARS-CoV-2 can also enter monocyte-derived macrophages and alveolar macrophages via phagocytosis, viral replication is restricted and there are low amounts of viral protein expression and virion production^30^. Notably, limited infection can still enhance pro-inflammatory cytokine expression in these macrophages^30^, highlighting macrophages as critical players in SARS-CoV-2 induced cytokine storm. To date, no studies have investigated the roles of individual SARS-CoV-2 proteins on macrophage immunity, underscoring an urgent need to understand how SARS-CoV-2 dysregulates macrophage functions to drive pathological hyperinflammation.

Here we demonstrate the varying roles of the SARS-CoV-2 N protein during macrophage infection. Specifically, we first show that the packaged N protein dampens the antiviral response, while *de novo* production of the N protein results in a hyperinflammatory phenotype during infection. To further unravel the pro-inflammatory role of the N protein, we generated THP-1 monocyte-derived macrophages that can inducibly express the SARS-CoV-2 Wuhan strain N protein. We systematically investigated the transcriptomic changes in macrophage innate immune signaling in response to endosomal (TLR7/8, TLR3) and cytoplasmic (RIG-I/MDA5) pattern recognition receptor stimulation in the presence and absence of the N protein. This revealed that the N protein specifically hyperactivates TLR-mediated inflammation while also dampening RIG-I mediated IFN activation. We then investigated these changes across SARS-CoV-2 variants and various coronaviruses, which revealed the delta variant N protein to be significantly more inflammatory than the omicron variant N protein. We also identified differential interactions with stress granule proteins between the delta and omicron variant N proteins that may drive hyperinflammation. Finally, we demonstrated the consequences of this hyperactivation by treating endothelial cells from the brain and heart with the supernatants of macrophages expressing these N proteins. Importantly, supernatant from TLR7/8-stimulated macrophages expressing the delta variant N protein significantly disrupts these barrier cells, correlating with key COVID-19 complications including cognitive impairment and cardiac injury. Overall, our study pinpoints the SARS-CoV-2 N protein as a major driver of macrophage dysregulation leading to severe COVID-19.

## RESULTS

### Incoming and *de novo*-expressed nucleocapsid elicit opposing immune responses in macrophages

Macrophages play a major role in SARS-CoV-2-associated complications and immunopathogenesis^31^; however, whether they are directly infected or indirectly manipulated by free SARS-CoV-2 proteins remains controversial. Compared to lung resident macrophages, THP-1 monocyte-derived macrophages are minimally infected by SARS-CoV-2. However, this can be overcome by overexpressing both CD169 and ACE2 receptors, resulting in productive SARS-CoV-2 infection^30^. While direct infection will result in a multitude of effects on macrophages, engulfment and clearance of SARS-CoV-2 infected apoptotic cells may also result in SARS-CoV-2-mediated macrophage manipulation^26,29,30,32^. It is known that the SARS-CoV-2 N protein is a master manipulator of the innate immune response across various cell types. To this end, we demonstrate that N-expressing HEK293T cells have dampened IFN activation and NF-κB signaling **(Supplemental Fig. S1A and S1B)**. Importantly, these data agree with previous studies that showed N as an antiviral antagonist in various cell types^16,33^. Since macrophages can encounter the N protein through many routes, and the exact effects of the N protein remain unclear, we first sought to determine which sources affect macrophages and to what extent/direction.

We first sought to understand N-mediated effects on macrophage activation during the early stages of infection. We mimicked the engulfment process by delivering the N protein into THP-1-derived macrophages with non-replicative SARS-CoV-2 virus-like particles (VLPs)^34^. We did this specifically in macrophages with or without overexpression of specific receptors (CD169, or both CD169 and ACE2). We then compared representative cytokine and chemokine expression of these macrophages after 24 hours of VLP incubation with an additional 4-hour treatment with a single-stranded RNA-mimic (CL097) that stimulates TLR7/8 **(Fig. 1A)**. It is interesting that the THP-1 macrophages overexpressing CD169 and CD169 with ACE2 are more responsive to CL097-mediated TLR7/8 stimulation, compared to parental THP-1 macrophages, potentially due to enhanced phagocytic activity after CD169 overexpression. Overall, VLP treatment followed by CL097 stimulation resulted in decreased expression of all genes tested in the parental macrophages relative to no VLP treatment; however, only CCL4 and IL-8 were statistically significant **(Fig. 1D and 1F)**. Meanwhile, the same trend was observed for the macrophages overexpressing CD169 and CD169 with ACE2, where all genes tested were significantly suppressed with VLP treatment, with the exception of IL-8 **(Fig. 1B-F)**. These data indicate that the incoming N protein acts as an antiviral antagonist, likely promoting a favorable environment for viral replication.

**Figure 1.**
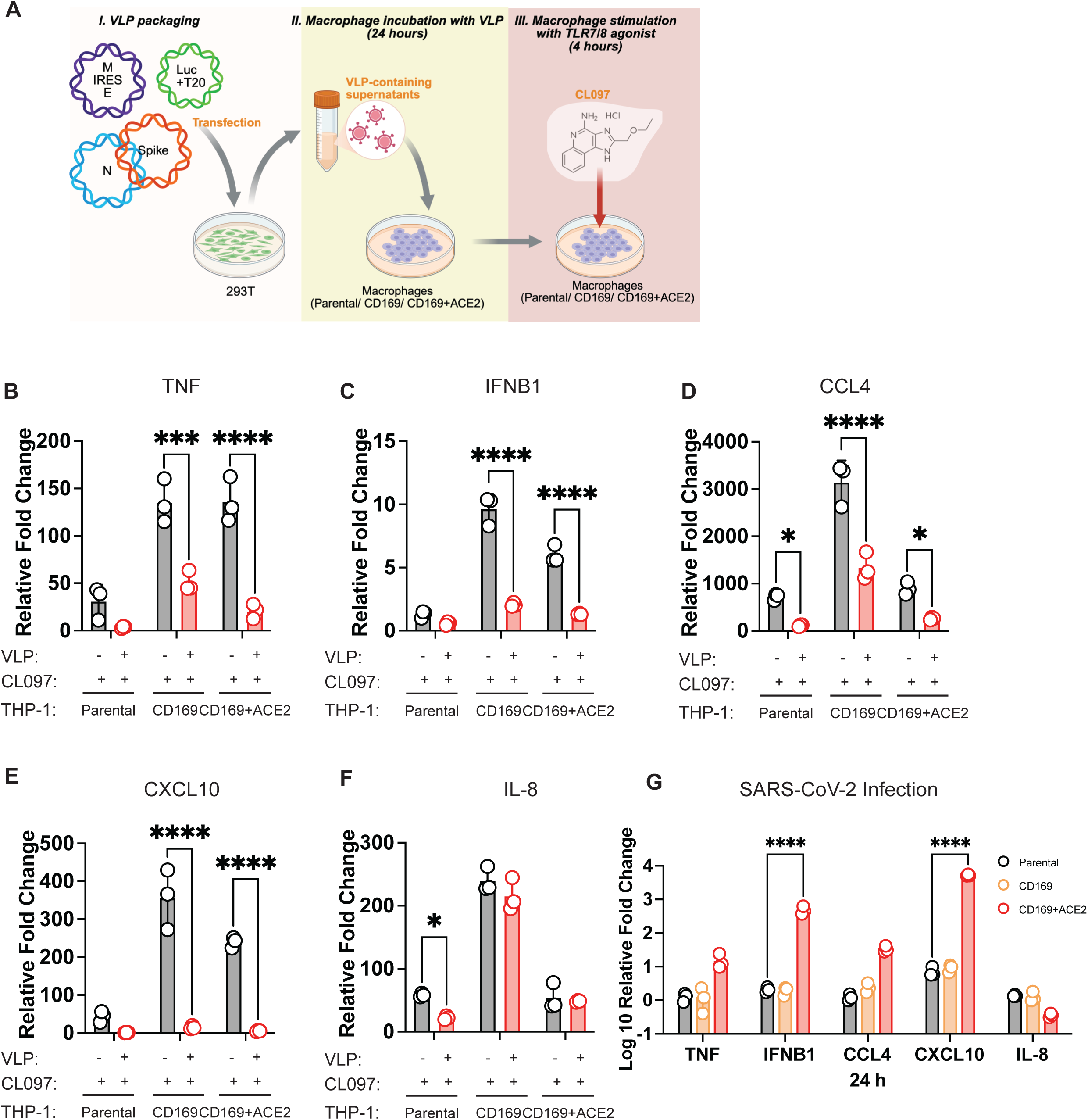
The effect of incoming or *de novo*-synthesized N protein on the induction of pro-inflammatory cytokines and chemokines in macrophages. **(A).** The schematic showing the SARS-CoV-2 VLP packaging and VLP incubation followed with TLR7/8 stimulation on different macrophage cell lines. **(B-F).** The VLP regulations on TLR7/8 stimulation-induced cytokine and chemokine expressions. The parental THP-1 macrophages or THP-1 macrophages overexpressing corresponding receptors (CD169 or CD169+ACE2) were incubated with VLPs (Wuhan strain structural proteins) for 24 hours. The macrophages were then stimulated with CL097 for 4 hours. The relative fold changes of the expressions of representative cytokines and chemokine (TNF, IFNB1, CCL4, CXCL10 and IL-8) are evaluated through RT-qPCR, by comparing to untreated cells. Data representative of two independent experiments. Asterisks indicate statistically significant fold change difference of CL097-treated cells vs VLP+CL097-treated cells (Two-way ANOVA and Šidák’s multiple comparisons test: *, p<0.05; ***, p<0.001; ****, p<0.0001). **(G).** The representative cytokine expressions induced by SARS-CoV-2 infections in THP-1 macrophages, CD169-overexpressing THP-1 macrophages and CD169+ACE2 overexpressing macrophages. The parental THP-1 macrophages or THP-1 macrophages overexpressing corresponding receptors (CD169 or CD169+ACE2) were inoculated with SARS-CoV-2 of MOI 1 and infected for 24 hours. The expressions of TNF, CCL4, CXCL10, IL-8 and IFNB1 are normalized and quantified by RT-qPCR detection. The relative fold changes are evaluated by the comparisons to uninfected cells and plotted into bar graphs in Log_10_ scale. Data representative of two independent experiments. Asterisks indicate statistically significant fold change differences of SARS-CoV-2-infected CD169 THP-1 macrophages and CD169+ACE2 THP-1 macrophages as compared to SARS-CoV-2-infected parental macrophages (Two-way ANOVA and Dunnett’s multiple comparisons test: ****, p<0.0001).

We next interrogated how *de novo* synthesis of the N protein during infection affects macrophages. Specifically, we infected the same macrophage lines with the parental strain of SARS-CoV-2 and measured the effects on the same cytokines and chemokines **(Fig. 1G)**. We found that the SARS-CoV-2 infection of THP-1 macrophages overexpressing CD169 with ACE2 upregulates the expression of most cytokines and chemokines, except for IL-8 which is downregulated. The expression levels of IFNB1 and CXCL10 increase dramatically, by approximately 200-fold and 5000-fold, respectively. The infection results suggest that new viral protein synthesis and multi-round infection promotes inflammation, in contrast to packaged N protein. Overall, these data show distinct roles for the N protein that aim to restructure the cellular environment to favor viral replication versus spread in a temporal manner, potentially contributing to the shift from early immune evasion to later hyperinflammation in COVID-19.

### Nucleocapsid enhances TLR-driven inflammation and suppresses RLR-mediated antiviral signaling

We have shown that the newly synthesized SARS-CoV-2 N protein can be pro-inflammatory during viral replication cycle. To further investigate and break down the pathways by which the N protein mediates this phenotype, we constructed THP-1 monocyte-derived macrophage cell lines with dox-inducible expression of the Wuhan strain N protein using lentivirus transduction. Given that SARS-CoV-2 is an RNA virus, it can activate endosomal (TLR7/8, TLR3) or cytosolic (RIG-I/MDA5) RNA sensors. We therefore treated these macrophage cell lines with molecules to stimulate these PRRs during N protein expression prior to bulk RNA-sequencing (RNAseq). Specifically, we stimulated TLR7/8 or TLR3 using free single-stranded RNA (CL097) or double-stranded RNA (poly(I:C)), respectively, and activated RIG-I/MDA5 by transfecting cells with poly(I:C). Multidimensional scaling (MDS) plots show that untreated parental and N-expressing macrophages cluster together **(Fig. S2A)**. While RIG-I/MDA5-stimulated N-expressing cells cluster near parental controls, TLR-stimulated N-expressing cells shift far from their parental controls **(red and green arrows, Fig. S2A)**. These data indicate that the N protein selectively drives large transcriptional changes in TLR pathways.

For each PRR stimulation, our RNAseq analysis identified differentially expressed genes (DEGs; False Discovery Rate (FDR)<0.05) that fit into 3 categories: 1) DEGs unique to N-expressing macrophages, 2) DEGs unique to parental macrophages, or 3) DEGs shared by N-expressing and parental macrophages **(Fig. 2A)**. As predicted by our MDS plots, the N protein dramatically alters the number of genes that are significantly regulated by TLR stimulation (TLR7/8 stimulation (left): 1422 DEGs unique to N-expressing macrophages vs 472 DEGs unique to parental macrophages; TLR3 stimulation (middle): 2827 DEGs unique to N-expressing macrophages vs 520 DEGs unique to parental macrophages). Interestingly, RIG-I/MDA5 stimulation (right) triggers the largest gene expression changes in macrophages, though a majority of these DEGs are shared, consistent with our MDS plots (1979 DEGs unique to N-expressing macrophages vs 1130 DEGs unique to parental macrophages; 3691 shared DEGs).

**Figure 2.**
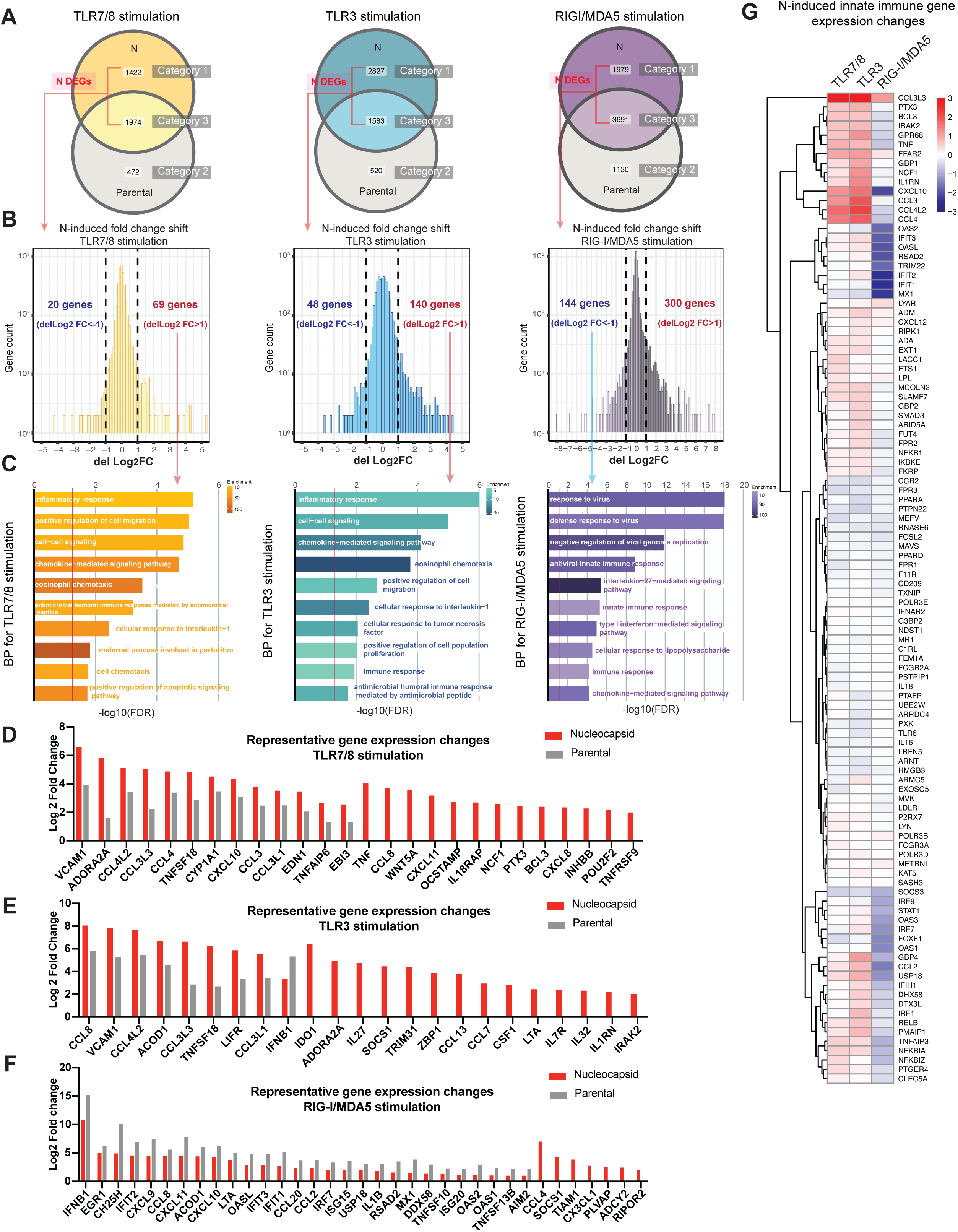
Nucleocapsid enhances TLR-driven inflammation and suppresses RLR-mediated antiviral signaling. **(A).** The Venn diagrams of differentially expressed genes (DEGs) in parental macrophages and N-expressing macrophages after PRR stimulations. The gene expression fold changes induced by TLR7/8, TLR3, RIG-I/MDA5 stimulation are compared in either parental macrophages or N-expressing macrophages. The genes are considered as DEGs with a filtration of FDR <0.05, regardless of values of Log2 Fold Changes (PRR-stimulated macrophages vs untreated macrophages). Overall, the DEGs are classified into 3 categories: 1) DEGs unique to N-expressing macrophages, 2) DEGs unique to parental macrophages, or 3) DEGs shared by N-expressing and parental macrophages. CL097 incubation (TLR7/8 stimulation) induced significant expression changes of 1974 genes in both N-expressing macrophages and parental macrophages, and 1422 genes only in N-expressing macrophages. Poly(I:C) incubation (TLR3 stimulation) induced significant expression changes of 1583 genes in both N-expressing macrophages and parental macrophages, and 2827 genes only in N-expressing macrophages. Poly(I:C) transfection (RIG-I/MDA5 stimulation) induced significant expression changes of 3691 genes in both N-expressing macrophages and parental macrophages, and 1979 genes only in N-expressing macrophages. **(B).** The histograms of N-induced gene expression changes. The histograms from left to right show N influence on induction changes of DEGs in TLR7/8, TLR3 or RIG-I/MDA5 stimulation by evaluating the delLog_2_Fold Change distribution in each condition. To obtain delLog_2_FC, the N log_2_ Fold Change of DEGs (PRR-stimulated N-expressing macrophages vs unstimulated N-expressing macrophages) is compared to the WT log_2_ Fold Change of DEGs (PRR-stimulated parental macrophages vs unstimulated parental macrophages). **(C).** The biological process (BP) analysis of N-induced DEGs in TLR7/8, TLR3 and RIG-I/MDA5 stimulations. Left: Top 10 enriched gene ontology terms of 69 N-upregulated DEGs (delLog2FC>1) in macrophages after TLR7/8 stimulation. Middle: Top 10 enriched gene ontology terms of 140 N-upregulated DEGs (delLog2FC>1) in macrophages after TLR3 stimulation. Right: Top 10 enriched gene ontology terms of 144 N-downregulated DEGs (delLog2FC<1) in macrophages after RIG-I/MDA5 stimulation. The bars exceeding -log10(FDR) cutoff (red line, FDR=0.05) shows significant enrichments. **(D-F).** Fold change bar graphs of representative DEGs dramatically regulated by N in PRR-stimulated macrophages. Log_2_ fold changes of DEGs exhibiting enhanced expression in both parental and N-expressing macrophages to different extents, and those selectively upregulated in N-expressing macrophages after TLR7/8 stimulation **(D)**, TLR3 stimulation **(E)** or RIG-I/MDA5 stimulation **(F)**. **(G).** Heatmap of DEGs associated with the innate immune response, showing N-induced expression changes across all PRR stimulation conditions. The DEGs in parental macrophages and N-expressing macrophages in all PRR-stimulated conditions are selected and filtered with innate immune response-related BP and KEGG pathway terms (GO:0140374, GO:0051607, GO:0009615, GO:0032728, GO:0070106, GO:0006954, GO:0045071, GO:0050729, GO:0045087, GO:0060337, GO:0032760, GO:0050728). The innate immune response-related DEGs are then plotted into heatmap colored with delLog_2_ Fold Changes of different PRR stimulation. The delLog2 Fold Changes are calculated as described in Figure 2B which are the comparison of N log_2_ Fold Change of DEGs to WT log_2_ Fold Change of DEGs under the same PRR stimulation condition. Red: DEGs upregulated by N after PRR stimulation. Blue: DEGs downregulated by N after PRR stimulation.

Next, we focused on the DEGs that fit into categories 1 and 3, which include DEGs unique to N-expressing macrophages and shared DEGs (annotated as **N DEGs in Fig. 2A**). It is notable that N expression can either further enhance or suppress the expression of these DEGs to varying degrees. To quantitatively determine the directionality and magnitude underlying N-mediated changes in gene expression, we calculated the difference in fold changes (delLog_2_Fold Change; delLog_2_FC) for N DEGs between the N-expressing macrophages and parental macrophages for each PRR stimulation. We then plotted the delLog_2_FC distribution of N DEGs for TLR or RLR stimulation **(Fig. 2B)**. Interestingly, the histograms show that the delLog_2_FC values slightly distributed to the right across all the PRR stimulation conditions, although there are several genes that are also downregulated. We have thus far shown that the N protein strongly dysregulates TLR and RLR pathways through both enhancement and suppression of specific genes.

To further unravel the pathways affected by the N protein, we performed gene ontology (GO) analysis on all hyperactivated genes (delLog_2_ FC >1) and all suppressed genes (delLog_2_ FC <-1) across each receptor pathway **(Fig. 2C and S2B)**. However, due to the low number of suppressed genes for either TLR pathway, no GO terms were enriched and thus were not plotted. Strikingly, the 69 and 140 genes highly upregulated by N in TLR7/8 and TLR3 stimulation are significantly enriched in inflammation-related GO terms **(Fig. 2C, left and middle plots)**, highlighting the pro-inflammatory nature of the N protein on TLR pathways in macrophages. In contrast, the 300 genes that are hyperactivated in RIG-I/MDA5 stimulation do not significantly overrepresent any GO terms **(Fig. S2B)**. Notably, the 144 genes downregulated by N upon RIG-I/MDA5 stimulation are mostly related to the GO terms related to the antiviral response **(Fig. 2C, right plot)**. This suggests that the N protein promotes the inflammatory response upon TLR stimulation and antagonizes the host IFN response upon RLR stimulation.

We next wanted to directly compare the expression fold changes of representative N DEGs related to innate immune response between the parental and N-expressing macrophages under every PRR stimulation **(Fig. 2D-2F and Supplementary Table 1)**. In TLR7/8 and TLR3 stimulation, the presence of the N protein leads to enhanced expression of pro-inflammatory cytokine transcripts relative to parental macrophages. Specifically, genes involved in COVID-19 cytokine storm, including CCL4, CXCL10, TNF, CCL8, etc.^35^, are dramatically elevated (Log_2_FC>4) in N-expressing cells. Interestingly, IFNB1 is suppressed upon TLR3 stimulation in N-expressing macrophages. In contrast, the presence of the N protein generally suppresses expression of antiviral ISGs (including CH25H, OASL, IFIT1, etc.), cytokines and chemokines upon RIG-I/MDA5 stimulation. For example, both IFNB1 and CH25H are decreased by about 32-fold in the presence of the N protein relative to parental macrophages. These results demonstrate that N steers the TLR-mediated response to a more inflammatory direction and broadly dampens the RLR-mediated response.

Finally, we wanted to compare the effects of the N protein across all three stimulus conditions. Therefore, we took 729 DEGs shared among all 6 experimental groups (TLR7/8, TLR3, or RIG-I/MDA5 stimulated N-expressing or parental macrophages) **(Supplementary Table 2)**, and generated a heatmap with 106 genes that are enriched in inflammation and IFN pathways based on the GO terms in BioMart database^36^ **(Fig. 2G)**. The innate immune genes clustered at the top and bottom of the heatmap demonstrate contrasting regulatory roles of the N protein in TLR7/8/3 and RIG-I/MDA5 pathways. For example, inflammatory innate immune genes are significantly upregulated by N in the TLR pathways but downregulated in the RIG-I/MDA5 pathways. These include inflammatory cytokines and chemokines such as TNF, CCL3, CCL4, CCL4L2, and CXCL10 **(Fig. 2G, top)** and components of the NF-κB pathway such as IRAK2, BCL3, RELB, TNFAIP3, and NFKBIA **(Fig. 2G, top and bottom)**. In contrast, the IFN pathway components and negative regulators such as STAT1, IRF7, IRF9, and USP18 **(Fig. 2G, bottom)** and antiviral ISGs such as OASL, IFIT1, IFIT2, and IFIT3 **(Fig. 2G, top)** are dramatically suppressed in RIG-I/MDA5 pathway **(Fig. 2G, bottom)**. Interestingly, CCL3L3, an inflammatory chemokine, and FFAR2, a fatty acid receptor involved in inflammatory response, are both enhanced by N in all three PRR pathways. Together, these data emphasize the role of the N protein as a master manipulator of the cellular environment, which promotes the inflammatory response and dampens the antiviral response in a stimulus-specific manner.

### The delta variant N protein is highly pro-inflammatory regardless of stimulation

Since we observed strong pro-inflammatory gene signatures in the TLR7/8 and TLR3 stimulated macrophages with N overexpression, we wanted to identify the most common downstream transcription factors that are disrupted by the N protein in these pathways. We predicted the most relevant transcription factors by mapping 69 upregulated genes in TLR7/8 stimulation and 140 upregulated genes in TLR3 stimulation into the TRRUST database^37,38^ **(Fig. 3A and 3B)**. The analysis clearly demonstrated that RELA and NFKB1 rank as top transcription factors that regulate the N-induced genes in both TLR7/8 and TLR3 pathways. The NF-κB subunits p50 and p65, encoded by NFKB1 and RELA, respectively, can form a dimer and subsequently translocate into the nucleus to enhance inflammatory gene expression. This is known as the classical NF-κB pathway initiated by TLR or TNF receptor stimulation^39^. In addition, other inflammatory response related transcription factors, including CEBPD and JUN, also exhibit high likelihood to be enhanced by the N protein in TLR7/8 stimulation. Although STAT1 is significantly enriched in both TLR stimulation, there are only a few genes (CXCL10, EDN1, and CCL3 in TLR7/8 stimulation; CXCL10, SOCS1, and IL27 in TLR3 stimulation) that map to the STAT1 network while most of the genes are overlapped with the NFKB1 and RELA networks. Overall, the transcription factor prediction highlights that the N protein specifically promotes the NF-κB-mediated inflammatory responses through TLR7/8 and TLR3 activation.

**Figure 3.**
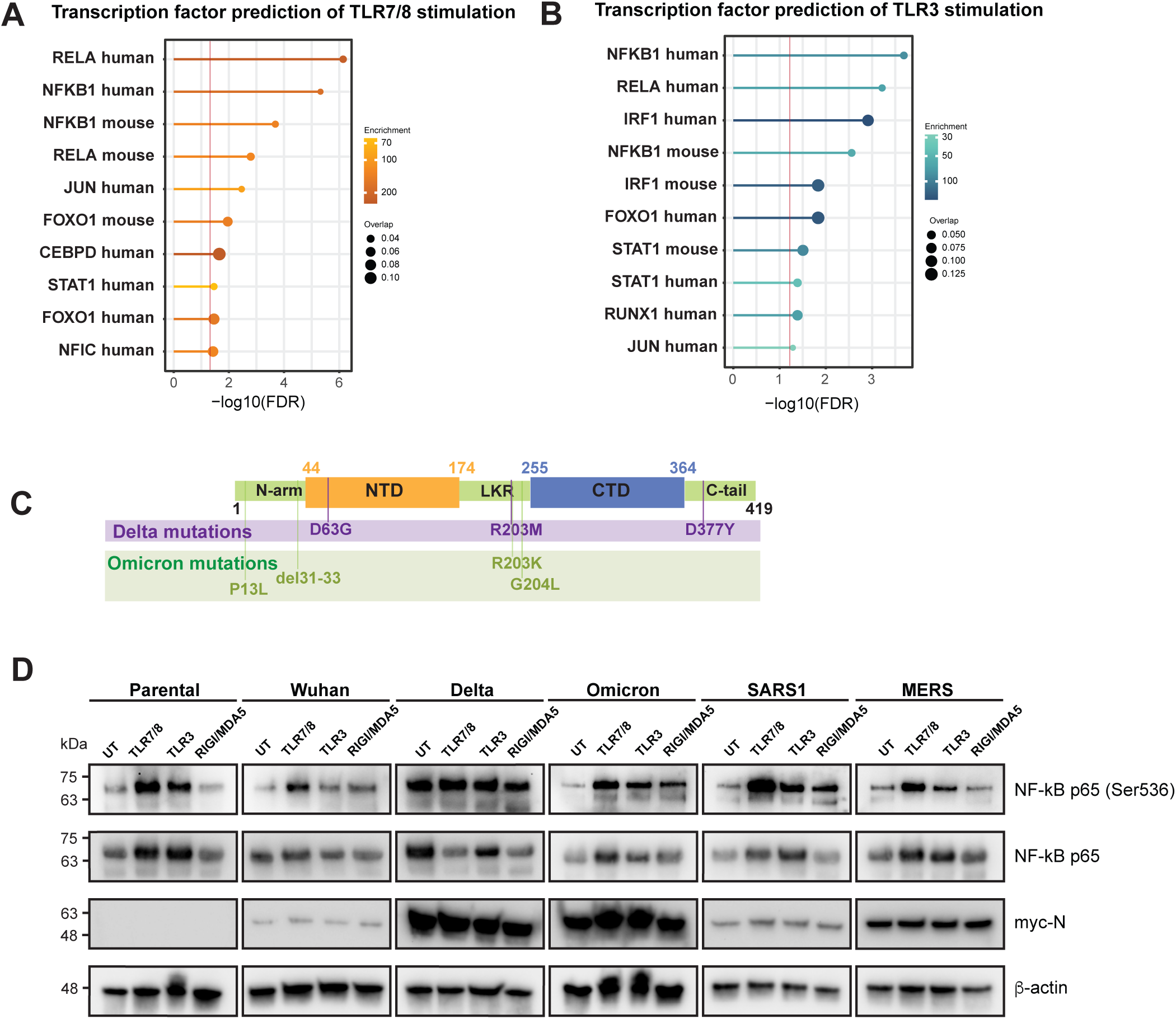
The delta variant N protein is highly pro-inflammatory regardless of stimulation. **(A-B).** The transcription factor predictions of N-mediated gene upregulations in TLR7/8 and TLR3 pathways. The 69 N-upregulated genes (Fig. 2B left, genes with delLog2 FC >1) after TLR7/8 stimulation and 140 N-upregulated genes (Fig. 2B middle, genes with delLog2 FC >1) after TLR3 stimulation are analyzed for responsible transcription factors through TRRUST, a manually curated database of human and mouse transcriptional regulatory networks. The top 10 predicted transcription factors based of enrichment significance (FDR) are listed. The gradient colors correspond to combined enrichment scores. The size of the dots corresponds to the overlap ratio of the N-upregulated genes in transcription factor-target genes in the database. The line segments exceeding -log10(FDR) cutoff (red line, FDR=0.05) shows significant enrichments. **(C).** The Mutation sites of delta and omicron variants in corresponding N domains. The domains of N protein can be divided into intrinsically disordered regions (IDRs) and conserved structural regions. The conserved structural regions consist of N terminal domain (NTD) and C terminal domain (CTD). The NTD and CTD are connected by an IDR, named linker region (LKR), and flanked by the other two IDRs, named N-arm and C-tail. The delta variant has D63G mutation in its NTD, R203M mutation in its LKR and D377Y mutation in its C-tail. The omicron variant has P13L and a three amino acid deletion (aa31-aa33) in its N-arm, R203K and G204L in its LKR. **(D).** The phosphorylation of NF-kB modulated by different coronavirus N proteins after PRR stimulations. The dox-inducible THP-1 derived macrophage cell lines that express different N proteins were treated with dox (2 mg/ml) for 48 hours to stimulate the N protein expressions of SARS-CoV-2 Wuhan strain, SARS-CoV2 delta variant, SARS-CoV-2 omicron variant, SARS-CoV-1, MERS. The parental THP-1 macrophages were also treated with dox(2 mg/ml) for 48 hours to serve as a control. The cells were sequentially stimulated with different PRR agonists, including CL097 incubation for TLR7/8 activation, poly (I:C) incubation for TLR3 stimulation, poly (I:C) transfection for RIG-I/MDA5 stimulation, for 4 hours. The cell lysates were then collected to detect the phosphorylation at Ser 536 in NF-kB p65 subunit by western blot.

Since the phosphorylation of NF-κB p65 at serine 536 is essential for its nuclear translocation and transcriptional activation^40^, we next investigated whether the N protein affects this modification. Specifically, we interrogated p65 phosphorylation under basal and all three stimulus conditions. Moreover, we were also interested in determining whether the effects of the N protein were consistent among the SARS-CoV-2 delta and omicron variants, as well as those of other betacoronaviruses, including SARS-CoV-1 and MERS-CoV. THP-1 monocytes with doxycycline-inducible stable expression of the N proteins of the SARS-CoV-2 delta and omicron variants, SARS-CoV-1, and MERS-CoV were constructed through lentivirus transduction. We generated the variant N proteins by introducing the following nonsynonymous mutations: D63G, R203M, and D377Y (for delta), and P13L, R203K, G204L, and deletion of residues 31-33 (for omicron) **(Fig. 3C)**. Interestingly, the specific mutations for the delta and omicron variants dramatically increase the expression of the N protein in THP-1 derived macrophages **(Fig. S3A)**.

We then stimulated the TLR7/8, TLR3 or RIG-I/MDA5 pathway in THP-1 derived macrophages expressing the N proteins of different coronavirus strains, variants, and species (Wuhan strain, delta variant, omicron variant, SARS-CoV-1, MERS), and measured p65 phosphorylation at serine 536 **(Fig. 3D)**. We found an increase of NF-kB p65 phosphorylation in parental macrophages upon TLR stimulation, while no difference between untreated and RIG-I/MDA5 stimulation was observed. Meanwhile, the N proteins from SARS-CoV-2 delta and omicron variants, and SARS-CoV-1 enhance p65 phosphorylation than N from Wuhan strain after PRR stimulation, especially for the RIG-I/MDA5 pathway. Strikingly, the expression of the delta variant N protein alone can hyperactivate p65 phosphorylation at the basal level, with no need for any PRR stimulation **(Fig. 3D)**. This suggests that the N protein from the delta variant is highly inflammatory. Overall, these data suggest that the pro-inflammatory nature of the N protein is mostly conserved among pathogenic coronaviruses.

The transcriptomics results **(Fig. 2C-2G)** clearly demonstrated the inhibition of ISGs by the Wuhan strain N protein upon RIG-I/MDA5 stimulation. Given that the phosphorylation of STAT1 at tyrosine 701 is crucial for initiating ISG expression, we probed for STAT1 phosphorylation in PRR-stimulated macrophages expressing the various N proteins previously tested **(Fig. S3B)**. Only TLR3 and RIG-I/MDA5 stimulation led to STAT1 phosphorylation in parental and Wuhan strain N-expressing macrophages. Interestingly, the N proteins of the delta and omicron variants show greater suppression of STAT1 phosphorylation than that of the Wuhan strain in the RIG-I/MDA5 pathway. The N proteins of SARS-CoV-1 and MERS-CoV also dramatically suppress STAT1 phosphorylation after RLR stimulation. In contrast, the N proteins of the SARS-CoV-2 Wuhan strain, and delta and omicron variants, and MERS-CoV further increase STAT1 phosphorylation mediated by TLR3 activation. This is consistent with the RNA-seq results where N upregulates some of the ISGs in the TLR3 pathway **(Fig. 2G, bottom)**, including GBP4, USP18, DHX58, DTX3L, etc. These findings indicate that coronavirus N proteins broadly antagonize RLR-driven ISG induction by blocking STAT1 activation, while selectively enhancing STAT1 phosphorylation and ISG expression in the TLR3 pathway.

### The interaction with cGAS and stress granule proteins may contribute to the pro-inflammatory features of macrophages during SARS-CoV-2 infection

Our data thus far has demonstrated a clear, pro-inflammatory role for the SARS-CoV-2 N protein, yet the interactions underlying this role is unclear. Moreover, we have showed that the delta variant N protein is highly pro-inflammatory, while the omicron variant N protein has a milder phenotype despite similar expression levels and sequence identity. Thus, we utilized affinity purification-mass spectrometry (AP-MS) to identify potential macrophage factors that differentially interact with these N protein variants to drive different phenotypes. We immunoprecipitated myc-tagged delta and omicron variant N proteins from doxycycline-inducible macrophage stable cell lines using anti-c-myc agarose beads, followed by identification of N binding partners via liquid chromatography-tandem mass spectrometry (LC-MS/MS) **(Fig. 4A)**. Macrophage stable cell lines without doxycycline treatment served as negative controls. We then analyzed interaction data through Significance Analysis of INTeractome (SAINT). High confidence interactors (Bayesian FDR ≤0.05 and average spectral count ≥11) were selected for fold change calculation. We then plotted the 39 host proteins that interact with at least one N protein into a protein-protein interaction network (PPI) **(Fig. 4B)** to visualize differences in interaction candidates.

**Figure 4.**
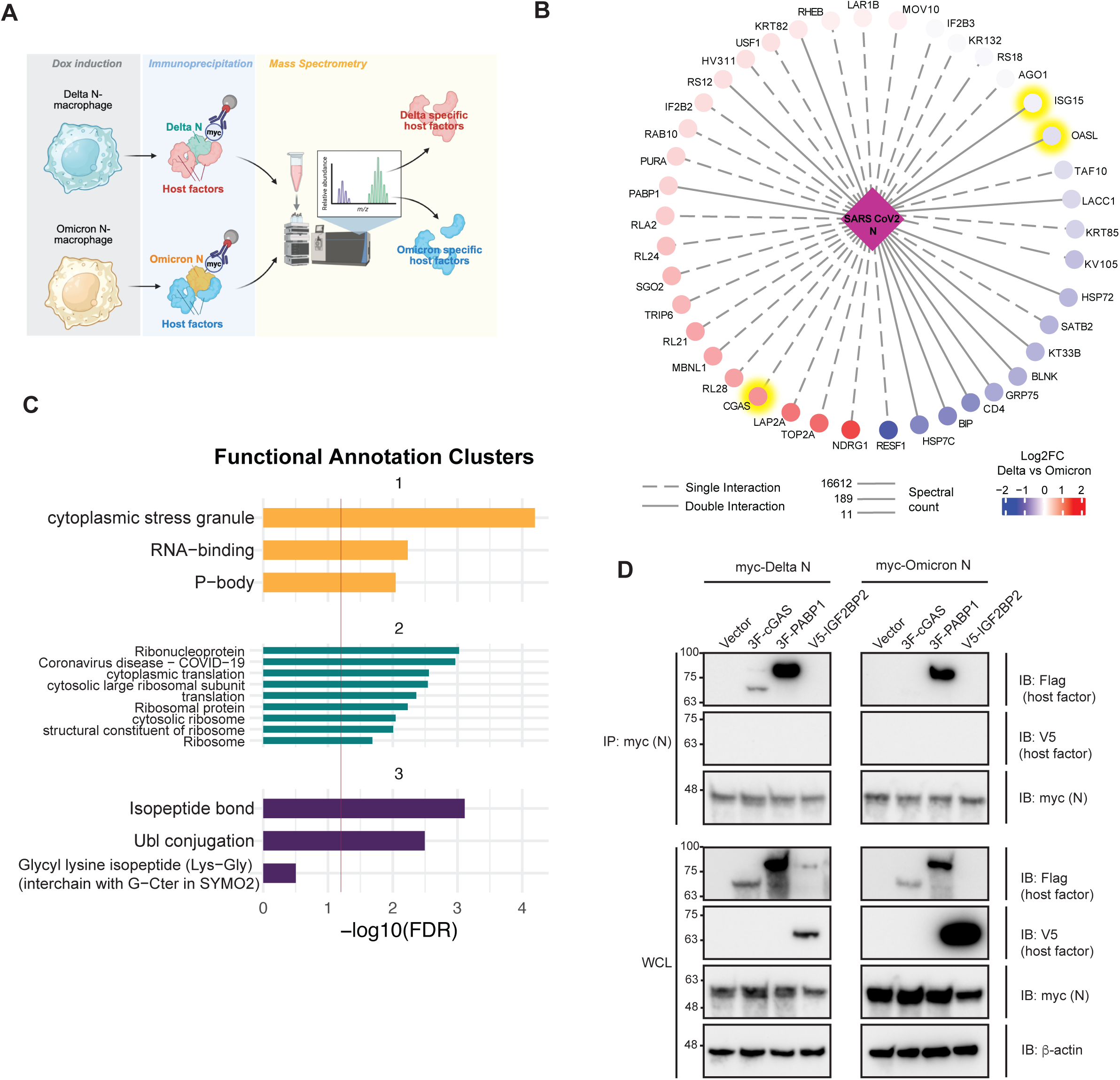
Identify the macrophage host factors that differentially interact with delta variant N and omicron variant N through AP-MS. **(A).** The schematic of AP-MS to identify specific host factors interacting with delta variant N and omicron variant N in macrophages. The corresponding dox-inducible THP-1 derived macrophages cell lines were either untreated or stimulated with dox (2 μg/ml) for 48 hours to express delta or omicron variant N. The delta or omicron N protein was immunoprecipitated with anti-myc antibodies conjugated on agarose beads through N-terminally fused myc tags. The host factors that were co-immunoprecipitated with the N proteins were later identified by LC-MS/MS. **(B).** Host factors that have confidential interactions with the N proteins of SARS-CoV2 variants. The 39 identified host proteins that pass the filtrations of SAINT BDFR≤ 0.05 and average spectral count ≥11 is calculated for their induction fold changes in delta N-expressing macrophages vs control macrophages (dox-untreated cells) or omicron N-expressing macrophages vs control macrophages (dox-untreated cells). The colors in the protein-protein interaction (PPI) network reflect the comparison of delta Log2 FC vs omicron Log2 FC, showing the interaction strength of the host proteins to different variants (Red: More interaction with delta variant N; Blue: More interaction with omicron variant N). The dashed line in PPI network shows that the host protein interacts at least one SARS-CoV2 variant N, while the solid line shows that the host protein interacts with both delta and omicron N. **(C).** The functional annotation clusters of confidential host interactors of N protein. The 39 host proteins in Fig. 4B were evaluated for their relationship to annotation terms in gene ontology and pathway analysis in DAVID database and grouped into annotation groups based on the Enrichment scores. The functional annotation terms are listed on the y-axis and the FDR of enrichment significance are ranged on the x-axis. The bars exceeding -log10(FDR) cutoff (red line, FDR=0.05) shows significant enrichments. **(D).** Co-immunoprecipitation validation of delta and omicron variant N protein interactors. HEK293T cells were transfected with plasmids expressing myc tagged delta or omicron variant N protein together with different host factors, including 3xFlag-cGAS (3F-cGAS), 3xFlag-PABP1 (3F-PABP1) and V5-IGF2BP2. Co-transfections of myc tagged delta/omicron variant N protein with empty vector are served as controls. Twenty-four hours post transfection, the cells were lysed and immunoprecipitated by anti-c-myc agarose beads. Immunoblot was probed to check for delta or omicron variant N binding to these host factors.

To appreciate the host biological functions that N proteins interfere with, we clustered the gene ontology and pathway annotations of the 39 host proteins through the DAVID database **(Fig. 4C and Supplementary table 3)**. Compared to the omicron variant, the delta variant N protein interacts more with a group of proteins such as MBNL1, PABP1, IGF2BP2 (gene name: IF2B2), LAR1B, MOV10, and IGF2BP3 (gene name: IF2B3) related to cytoplasmic stress granule formation **(Fig. 4B and Supplementary table 3)**. Interestingly, the delta variant N shows confident interactions with several ribosomal proteins (RL21, RLA2, RL24, RL28, RS12) suggesting the interference of the SARS-CoV2 N protein with host translation **(Fig. 4B and Supplementary table 3)**. Interestingly, nearly half of the N interactors are enriched in post-translational modification pathways of isopeptide bond and Ubl conjugation. It is known that isopeptide bonds are commonly formed during ubiquitination and SUMOylation, which implies that the N protein interactors may be involved in these processes. Although not ranked among the top three annotations, both delta and omicron variant N proteins significantly interfere with the host antiviral defense response (**Supplementary Table 3**, Enrichment Score: 1.42). Consistent with the PPI network analysis, the Omicron N protein displays stronger interactions with antiviral effectors ISG15 and OASL (highlighted in **Fig. 4B**). These results suggest that the SARS-CoV-2 N protein potentially influences gene expression in macrophages by modulating stress granule formation, host translation, and post-translational modification. Considering that stress granule formation can affect the expression of several components involved in the NF-κB pathway^41^, it is likely that the N protein regulates macrophage inflammation through interactions with stress granule components.

Notably, according to the PPI network, the delta variant N protein has a strong and specific interaction with the cytosolic DNA sensor cGAS to potentially trigger IFN and pro-inflammatory cytokine expression. Recent study revealed that cGAS undergoes condensation into stress granules through liquid-liquid phase separation (LLPS) to be primed for activation^42–44^. To confirm whether the delta variant N specifically interacts with cGAS and other stress granule components, we overexpressed myc-tagged delta or omicron variant N in HEK293T cells together with different host factors (cGAS, PABP1, or IGF2BP2 tagged with 3xFlag or V5). We pulled down the myc-tagged delta or omicron variant N protein and detected host factors that co-immunoprecipitated with the N protein **(Fig. 4D)**. We found that both delta and omicron variant N proteins have strong interactions with PABP1, while only the delta variant N protein interacts with cGAS. Neither delta nor omicron variant N protein binds to IGF2BP2, although this may be due to its low expression level in cells overexpressing delta variant N. These results strongly indicate that the delta variant N protein may regulate cGAS-mediated pro-inflammatory response through direct interaction or mediation through the stress granule.

### SARS-CoV-2 N amplifies cytokine secretion and triggers endothelial barrier breakdown

Having characterized N protein-driven pro-inflammatory transcriptional and signaling changes, we next examined the downstream effects on secreted factors and vascular integrity. Using a multiplex immunoassay, we determined the concentration of 71 different cytokines in the media of parental and either the SARS-CoV-2 Wuhan strain, delta variant, or omicron variant N-expressing macrophages with or without TLR7/8 stimulation **(Supplementary Table 4)**. Among these 71 cytokines and chemokines, we identified dysregulated secretion of 5 representative pro-inflammatory factors most of which are validated with remarkable induction by the Wuhan strain N protein at transcriptional levels in **Fig. 2D (Fig. 5A-E)**. We specifically show TNF, CCL4, and CXCL8 as they directly overlap with the transcriptomic data; however, we have also included CXCL9 and CXCL10 as they are closely related to and often correlate with CXCL11, which was also in the transcriptomic data. Among the unstimulated macrophages, the macrophages expressing the delta variant N protein are the most hyperactivated as there are higher amounts of TNF, CCL4 (not statistically significant), CXCL8, and CXCL10 relative to parental macrophages. Moreover, macrophages expressing the SARS-CoV-2 Wuhan strain or delta variant N protein are the most hyperactivated among the stimulated macrophages, with 4 or 5 out of 6 cytokines being secreted significantly more relative to parental macrophages, respectively. Interestingly, cytokine profiles for macrophages expressing the omicron variant N protein were the most unpredictable, as there were instances where the concentration of cytokine far exceeded that of others (CXCL8, unstimulated; **Fig. 5C**), and other instances where there was a strong decrease (TNF and CCL4, stimulated; **Fig. 5A-B**). Overall, the magnitude of cytokine induction varies by strain/variant and cytokine, with the delta variant N protein often inducing the most robust responses, while the omicron variant N protein elicited modest reduction in select cytokines.

**Figure 5.**
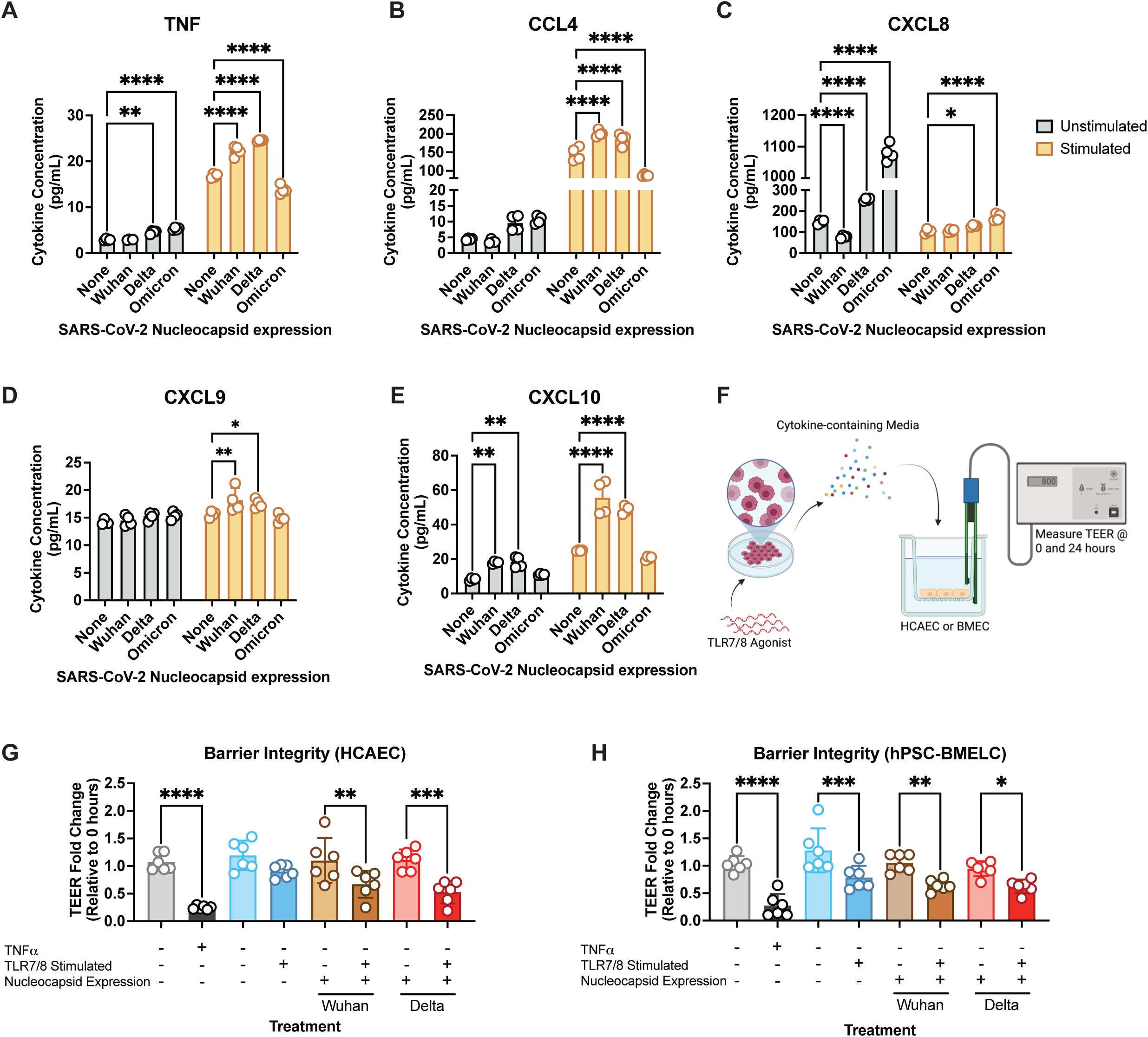
SARS-CoV-2 N amplifies cytokine secretion and triggers endothelial barrier breakdown. **(A–E)** Cytokine concentrations in the supernatant of parental THP-1–derived macrophages or those transduced with either SARS-CoV-2 nucleocapsid (N) from Wuhan, delta, or omicron variants, under unstimulated or TLR7/8-stimulated conditions. Cytokines were quantified using a 71-plex bead-based immunoassay (Eve Technologies). Representative cytokines shown include TNF (A), CCL4 (B), CXCL8 (C), CXCL9 (D), CXCL10 (E). **(F)** Schematic overview of the experimental setup for endothelial barrier assays. Conditioned media from the indicated macrophage treatments were applied to either human coronary artery endothelial cells (HCAECs) or stem cell-derived brain microvascular endothelial cells (hPSC-BMELCs). **(G–H)** Trans-endothelial electrical resistance (TEER) measurements in HCAECs (G) and hPSC-BMELCs (H) treated with conditioned media from the macrophage groups shown. Media from nucleocapsid-expressing, TLR7/8-stimulated macrophages significantly reduced TEER, indicating impaired barrier integrity. TNFα was included as a positive control for barrier disruption. All graphs show mean ± SD. Graphs H and I show combined data from two independent experiments. Asterisks indicate statistically significant differences (two-way ANOVA and Šidák’s multiple comparisons): *, p < 0.05; **, p < 0.01; ***, p < 0.001; ****, p < 0.0001. Non-significant differences are not shown.

To examine the functional impact of this cytokine-rich environment on vascular integrity, we transferred conditioned media from unstimulated and stimulated macrophages onto primary human coronary artery endothelial cells (HCAECs) and human pluripotent stem cell-derived brain microvascular endothelial-like cells (hPSC-BMELCs). Specifically, we focused on the effects from the SARS-CoV-2 Wuhan strain and delta variant N proteins since they were the most inflammatory from our cytokine profiling. We then measured transendothelial electrical resistance (TEER) as a proxy for barrier function (**Fig. 5F**) and found our positive control TNF-α to dramatically compromise the integrity of both endothelial cell barriers (**Fig. 5G-H**). When investigating changes in HCAEC integrity, we found that media from stimulated parental macrophages has no significant effect on HCAEC integrity **(Fig. 5G)**; however, media from either the Wuhan strain or delta variant N-expressing macrophages significantly reduced the barrier integrity down to 65% (Wuhan) or 50% (delta) of the original value. For hPSC-BMELCs, media from all stimulated macrophage cell lines resulted in a decrease in barrier integrity (**Fig. 5H**). Importantly, the changes in integrity were different: media from stimulated parental macrophages resulted in a decrease to 80%, while media from both the Wuhan strain and delta variant N-expressing macrophages reduced the original values down to about 60% of the original value. Together, these findings demonstrate that the SARS-CoV-2 N protein enhances cytokine secretion in a strain/variant-specific manner and promotes vascular leakage through paracrine inflammatory signaling.

## DISCUSSION

SARS-CoV-2, like many pathogenic viruses, has several immune antagonists that suppress host antiviral defenses to enable viral replication and spread. Multiple studies have identified several SARS-CoV-2 proteins—most notably non-structural protein (nsp) 1, nsp3, and ORF6—as potent IFN antagonists^15–17^. The N protein, though primarily recognized for its role in genome packaging, has also been implicated in blunting type I IFN response, particularly in the context of RIG-I activation in immortalized cell lines like HEK293T^8,15–17^. Our initial work recapitulates these findings, showing that N expression in HEK293T cells suppresses antiviral signaling downstream of RIG-I. To better reflect the physiological interactions in an immune-relevant system during infection, we investigated the activation of several PRRs in THP-1 monocyte-derived macrophages. Through this work, we identified a pro-inflammatory role for the N protein in a TLR-stimulated macrophage context. Moreover, we demonstrated that the N protein from various coronavirus strains and variants hyperactivate NF-kB, though the magnitude of this hyperactivation varies, with the delta variant having the most pro-inflammatory effect. Through proteomic analysis, we identified cGAS and several stress granule related proteins to interact more strongly with the delta variant N protein, which may contribute to its hyperinflammatory role. Finally, we demonstrated that the N protein leads to significantly more cytokine production, and as a result, induces more vascular leakage. Our work thus far establishes a novel and surprising function of the N protein, where it may interfere with cGAS and stress granule formation to trigger a hyperactive inflammatory response that contributes to cytokine storm and ARDS.

The N protein is the most abundantly expressed SARS-CoV-2 protein. In addition to its inhibitory role in innate immune responses, several clinical studies also revealed its pro-inflammatory role in facilitating disease severity^45,46^. Our RNA-seq data revealed that the N protein preferentially suppresses the expression of ISGs upon RIG-I/MDA5 stimulation, indicating that N-mediated IFN antagonism is unique to canonical RLR signaling. While this result supports existing literature, we also found that the N protein amplifies expression of inflammation-related transcripts upon TLR stimulation, suggesting that the N protein selectively manipulates immune signaling depending on the pathway engaged. These findings raise important questions regarding the mechanism of N-mediated immune modulation and the roles of the N protein in pathogenesis. Specifically, whether the N protein modulates similar pathways from HEK293T cells like TRIM25-mediated RIG-I ubiquitination in macrophages, or if alternative, macrophage-specific pathways are involved^8,23^. Moreover, while it is intuitive for pathogenic viruses to antagonize antiviral responses, it is less understood as to why pushing macrophages into a more pro-inflammatory state is evolutionarily advantageous. Future studies should explore whether this immune modulation is cell-type dependent and ultimately promotes SARS-CoV-2 pathogenesis.

Macrophages not only serve as key innate immune sensors but also play a central role in the hyperinflammatory responses associated with severe COVID-19^31^. Mononuclear phagocytes (MNPs) comprise 80% of the cells in bronchoalveolar lavage fluid from COVID-19 patients, and several of these MNPs are inflammatory monocyte-derived macrophages^28,47^; yet the mechanisms by which they are hyperactivated are unclear. Specifically, the field remains unsure as to whether indirect stimulation by peripheral SARS-CoV-2 infected cells, or if direct encounter with SARS-CoV-2 and interaction with viral immune modulators such as the N protein leads to chronic activation. One hypothesis is that delayed or chronic type I IFN signaling in COVID-19 contributes to the persistent, hyperactivated state of macrophages. One study showed that SARS-CoV-2 infection triggers TLR7 in plasmacytoid dendritic cells, leading to IFN-⍺ production that drives macrophages to a pro-inflammatory state^48^. An alternative hypothesis is that internalized SARS-CoV-2 results in macrophage-mediated hyperinflammation; however, whether SARS-CoV-2 can productively infect macrophages remains controversial. Ultimately, SARS-CoV-2 infection of macrophages largely depends on the expression of cellular receptors used for virus entry by various macrophage subtypes^26,30^. ACE2 is lowly expressed in macrophages, therefore SARS-CoV-2 harnesses lectin receptor-mediated phagocytosis to enter these cells. Siglec-1/CD169 mediates SARS-CoV-2 entry in monocyte-derived macrophages and alveolar macrophages, although with little to no production of infectious progenies^30^, while CD209 mediates virus entry in interstitial macrophages^26^. Therefore, we chose to investigate whether productive infection is required to induce transcriptional changes, or if packaged N protein in the virion is sufficient. Indeed, our data demonstrated that both mechanisms drive changes in macrophages, and that these changes are opposite in directionality. This suggests that the N protein has two roles in modulating the innate immune response during infection: 1) incoming N protein dampens the antiviral response to allow for effective replication, while 2) newly synthesized N protein may be more important for inducing a pro-inflammatory response that leads to further dissemination of the virus. Ongoing work will investigate whether these roles are temporally and spatially distinct, and whether blockade of newly synthesized N protein will prevent its pro-inflammatory functions.

With the continuing spread of SARS-CoV-2, we are witnessing viral evolution and human adaptation in real time as newer variants emerge and gain fitness advantages. Clinical studies showed a higher disease severity in individuals infected with the Wuhan strain and delta variant compared to the omicron variant. This was reflected by a greater magnitude of inflammatory cytokines and chemokines in the plasma induced by infection with the Wuhan strain and delta variant^49^. Indeed, introduction of the delta and omicron variant N proteins into our THP-1 monocyte-derived macrophage system leads to the discovery of several important differences. First, both the N proteins of the delta and omicron variants express significantly better compared to the Wuhan strain N protein despite all constructs being codon optimized. This suggests that these variants have developed more efficient expression in human cells. Next, relative to the Wuhan strain, the delta variant N protein is significantly more inflammatory, resulting in significant NF-κB phosphorylation across all conditions, and strikingly, under no stimulation. In line with clinical studies, the omicron variant N protein is less inflammatory, despite its similar expression as that of the delta variant N protein, suggesting intrinsic differences in the ability to promote inflammation. Ultimately, the mechanisms driving these different inflammatory profiles between variants remain elusive. Compared to the Wuhan strain, the N protein of the delta variant contains 3 mutations: D63G, R203M and D377Y, while the omicron variant has 4 mutations: P13L, deletion of 31-33, R203K and G204R. Interestingly, both the N proteins of the delta and omicron variants share a mutation at residue 203, but the delta variant has a more drastic change than the omicron variant. Additionally, it was reported that both the R203M and D377Y mutations in the N protein can outcompete RIG-I to bind to viral RNA, preventing IFN activation^24^. In the omicron variant, the R203K mutation together with G204R has been isolated as adaptive mutations associated with enhanced viral replication and infectivity^46,50^. Other studies suggested that post-translational modifications at K203 in the omicron variant N protein contributes to increased virulence through suppressing IFN expression^51^. While past studies have focused on the role of SARS-CoV-2 variant N proteins in IFN antagonism, future studies should investigate the roles of each mutation in promoting inflammation in macrophages.

While several studies have identified mechanisms of N-mediated dampening of the IFN response, it is not understood how the N protein may promote inflammation. To begin unraveling this mechanism, we performed comparative proteomics between the hyperinflammatory delta variant N protein and the less inflammatory omicron variant N protein. The proteomic results showed that the N protein of the delta variant interacts more strongly with stress granule components such as IGF2BP2/3, PABP1, and MOV10, and DNA sensor, cGAS. Stress granule formation is an important antiviral mechanism through host translation stalling which sequesters viral RNA transcripts and promotes innate immune response through condensation of cytosolic sensors, such as cGAS^43,44,52^. Current studies suggest that the N protein can be integrated into stress granules though LLPS and interfere with stress granule-mediated antiviral activity through interacting with critical components such as PKR, G3BP1, and MOV10^22,53^. Our immunoprecipitation results validated stronger interaction of the delta variant N protein with cGAS, which indicates variant-dependent effects on immune sensors associated with stress granules. It is reported that cGAS binds to G3BP1 and DDX3X to undergo condensation into stress granules for prompt activation^43,44^. Thus, the delta variant N protein which was found to interact more with stress granule components than the omicron variant N protein may hyperactivate cGAS and promote pro-inflammatory cytokine expression. Therefore, follow-up work will compare the impact of the N protein from different variants on stress granule components and associated proteins which can potentiate different inflammatory states.

Finally, we sought to better understand the clinical relevance of the hyperactive phenotypes we observed. We characterized cytokines secreted from TLR7/8-stimulated macrophages expressing different N proteins and found upregulation of several inflammatory factors. Among these, CCL4 (MIP-1β) stood out as most strongly correlating with cytokine signatures from COVID-19 patients^49^. CCL4 is a macrophage-secreted chemokine that recruits monocytes, T cells, and NK cells, and is consistently elevated in severe SARS-CoV-2 infection^49,54^. It is also capable of activating microglia and astrocytes and has been implicated in HIV-associated neurocognitive disorders, stroke-induced neuroinflammation, and blood-brain barrier leakage in COVID-19^54–57^. In line with these results, media from these hyperactivated macrophages greatly induced BMEC permeability *in vitro* compared to macrophages without the N protein. These properties strongly suggest a role for the N protein in COVID-19-associated neurological symptoms and long COVID brain fog, especially when it promotes secretion of these cytokines in excess upon innate immune activation. In parallel, we also observed HCAEC barrier disruption in our model, which has important implications for COVID-19-related cardiovascular pathology. HCAECs line the coronary arteries and are essential for maintaining vascular integrity and regulating immune cell trafficking^58^. Disruption of this barrier can promote myocarditis, atherosclerotic plaque destabilization, microvascular dysfunction, and thrombosis, particularly under pro-inflammatory conditions. In the context of COVID-19, such vascular complications have been widely reported^59^. Specifically, endothelial dysfunction, in addition to hyperinflammation, has been shown to contribute to disease severity and death in patients infected with SARS-CoV-2^60^. Our data therefore support a mechanistic model in which the N protein alone is sufficient to drive pathogenic cytokine responses in macrophages that in turn compromise critical endothelial barriers in the brain and heart. Together, these findings highlight N as a potent immune modulator with potential to exacerbate systemic complications of SARS-CoV-2 infection. Future studies should explore the feasibility of pharmacologically targeting N, or its downstream signaling pathways, to mitigate COVID-19-associated inflammatory damage.

Overall, this study reveals a previously unrecognized pro-inflammatory function of the SARS-CoV-2 N protein. We show that newly synthesized N protein, particularly from the Wuhan strain and delta variant, amplifies TLR-driven inflammation in macrophages. Proteomic analysis implicates N interactions with stress granule proteins and cGAS as a potential mechanism that promotes immune activation and inflammatory gene expression. These mechanisms drive excessive cytokine production in macrophages that disrupts endothelial integrity, pointing to a direct viral contribution to cytokine storm and vascular pathology. Together, these findings connect molecular events within infected cells to clinical outcomes such as brain and heart complications in severe COVID-19. Future therapeutic strategies that target N-mediated immune modulation could reduce the inflammatory burden of SARS-CoV-2.

## Supporting information

Table 5: Primers and antibodies

## Acknowledgements

We thank Ryan Kan from Dr. Aparna Bhaduri’s lab and Dr. Lifang Ye from Dr. Jeffrey Hsu’s lab at UCLA for their important support and expertise on establishing the HEK293T SARS-CoV-2 cell lines and HCAEC maintenance. We thank Dr. Rahm Gummuluru from Boston University for kindly providing us with CD169- and CD169+ACE2-expressing THP-1 cels. We thank Dr. LeAnn Nguyen, Dr. Serina Huang, Erin Kim, Martin Ruvalcaba, and Maria Villalba Nieto for their support and insights that helped this work move forward. We thank the James B. Pendleton Charitable Trust and the McCarthy Family Foundation for their generous support which provided critical equipment to the UCLA AIDS Institute. The following reagent was deposited by the Centers for Disease Control and Prevention and obtained through BEI Resources, NIAID, NIH: SARS-Related Coronavirus 2, Isolate hCoV-19/USA-WA1/2020, NR-52281. This work was funded in part by UCLA DGSOM – W.M. Keck Foundation Junior Faculty Award, Broad Stem Cell Research Center COVID-19 Research Award, JCCC-BSCRC Ablon Scholars Award, American Heart Association Mechanisms Underlying Cardiovascular Consequences Associated with COVID-19 and Long COVID Award to MMHL. ZY was supported by the Sydney Finegold Post-Doctoral Fellow Award. PAA was supported by the NIH under Award Number F31AI179235.

## MATERIALS AND METHODS

### Cell culture

THP-1 human monocytes were obtained from ATCC. The CD169-expressing THP-1 and CD169+ACE2-expressing THP-1 cell lines were kindly provided by Prof. Rahm Gummuluru at Boston University. The THP-1, CD169-expressing THP-1 and CD169+ACE2-expressing cell lines were cultured in Roswell Park Memorial Institute 1640 Medium (RPMI 1640, Gibco) supplemented with 10% FBS, 1X penicillin/streptomycin (P/S, Fisher Scientific), 1X non-essential amino acids (NEAA, Gibco), and 0.05 mM β-mercaptoethanol (Sigma-Aldrich). Human embryonic stem cells (H9; WA09, WiCell) were cultured on T 75cm^2^ flasks coated with 0.2mg/mL Matrigel (Corning) solution in 1:1 DMEM/Ham’s F12 (Gibco). Stem cells were grown with mTeSR1 (STEMCELL Technologies) with daily media changes. Human embryonic kidney (HEK) 293T cells were cultured in Dulbecco’s Modified Eagle Medium (DMEM, VWR) supplemented with 10% FBS. Human coronary artery endothelial cells (HCAECs) (Cell Applications) were cultured in Human Meso Endo Growth Medium (Cell Applications) on flasks coated with 0.1% gelatin solution (Millipore Sigma).

### THP-1 differentiation into macrophages

To differentiate THP-1, CD169-expressing THP-1 and CD169+ACE2-expressing THP-1 cells into macrophages, the cells were subjected to a 24-hour (h) stimulation with 50 ng/mL phorbol 12-myristate 13-acetate (PMA, Sigma-Aldrich) in RPMI 1640 supplemented with 10% human AB serum (Sigma-Aldrich), 1X NEAA, 1X P/S, which is followed by a 24 h rest in human-serum containing RPMI 1640.

### Human pluripotent stem cell-derived brain microvascular endothelial-like cell (hPSC-BMELC) differentiation and culture

Human pluripotent stem cells (hPSCs) were cultured and differentiated into hPSC-BMELCs as previously described^61^. Briefly, once hPSCs reached 70% confluency, mTeSR1 was replaced with 1:1 DMEM/Ham’s F-12 supplemented with 20% KnockOut Serum Replacer (Gibco), 1mM L-glutamine (Sigma Aldrich), 1X NEAA (Gibco), 0.1mM β-mercaptoethanol (Sigma Aldrich), 50ng/mL human basic fibroblast growth factor (STEMCELL Technologies). Cells were maintained using this medium for a minimum of five days with daily media changes. On the sixth day, medium was changed to Endothelial Cell Serum Free Media (EC SFM; Gibco) supplemented with 1X B-27 (Gibco), 10µM retinoic acid (Sigma Aldrich), and 20ng/mL human basic fibroblast growth factor (STEMCELL Technologies). Cells were maintained using this medium for a minimum of 48 hours. Differentiated cells were lifted off the culture vessel using StemPro Accutase (Gibco) and were washed several times prior to seeding at a density of 1×10^6^ cells/mL on plates coated with 400µg/mL type IV collagen from human placenta (Sigma Aldrich) and 100µg/mL fibronectin from human plasma (Millipore Sigma). hPSC-BMELCs were seeded in EC SFM supplemented with 1X B-27 and 10µM ROCK inhibitor and maintained in EC SFM supplemented with 1X B-27 with daily media changes.

### SARS-CoV-2 infection and virus-like particle (VLP) inoculation

The SARS-CoV-2 (Isolate hCoV-19/USA-WA1/2020, NR-52281, obtained through BEI Resources) stocks were prepared and titrated with Vero cells in a biosafety level 2+ lab. To infect the macrophages differentiated from THP-1 parental cells and stable cell lines that express CD169 or CD169+ACE2, virus stock was diluted in Dulbecco’s Phosphate Buffered Saline (DPBS) supplemented with 1% human AB serum (Sigma-Aldrich) and added to cells at a multiplicity of infection (MOI) of 2 plaque-forming units (PFU)/cell. The infection was carried out in triplicates in a 12-well plate with 5 × 10^5^ macrophages seeded per well. After 1 hour incubation with the virus, freshly made media (RPMI 1640 with 10% human serum) was added to cells. The total cellular RNAs were isolated for RT-qPCR detection.

To generate the SARS-CoV2 VLPs, we followed the protocol established by Syed and Taha^34^. Briefly, we transfected HEK293T cells in a 15-cm plate with CoV2-N (0.67, Addgene#177937), CoV2-M-IRES-E (0.33, Addgene#177938), CoV2-Spike-D614G (0.25, Addgene#177960), and Luc-PS9 (1, Addgene#177942) at the indicated mass ratios for a total of 40 μg of DNA. Total DNA and 120 μl PolyJet transfection reagent (SignaGen) was individually diluted in 1 mL serum free DMEM media. Diluted PolyJet was added to the diluted DNA and mixed gently. Transfection mixture was incubated for 15 min at room temperature and then added dropwise while gently swirling the plate. Media was replaced after 18 hours with 15 mL fresh DMEM containing bovine serum and penicillin/streptomycin. 42 hours after transfection, supernatant was collected and filtered with a 0.45 um pore size syringe filter. The macrophages at the amounts of 5 × 10^5^ cells/well were incubated with 540 μl VLP in 12-well plates for 24 hours, followed by a four-hour stimulation of CL097 in RPMI 1640 with 10% human serum at the concentration of 1.6 μg/ml.

### Construction of SARS-CoV-2 (Wuhan strain, delta and omicron variants), SARS-CoV-1, MERS-CoV N-expressing plasmids and tagged host factor (cGAS and PABP1) plasmids

All the primers that are used in gateway cloning, NEBuilder HiFi DNA Assembly, site-directed mutagenesis are listed in **Supplementary Table 5.**

The lentivirus plasmid expressing myc-tagged SARS-CoV-2 N (Wuhan strain) was constructed through Gateway cloning. Briefly, the myc-tagged Wuhan strain N protein flanked with attB sequences was codon-optimized and synthesized by Codex DNA Inc. The myc-N (Wuhan strain) fragment was first moved into pDONR221 (Invitrogen) vector through BP reaction by using Gateway™ BP Clonase™ II Enzyme mix, then transferred from pDONR221 to pCW57.1 (Addgene#41393) tetracycline (tet)-inducible lentivirus vector through LR reaction by using Gateway™ LR Clonase™ II Enzyme mix according to manufacturer’s instructions to generate pCW57.1-SARS-CoV-2 N (Wuhan strain).

To generate lentivirus plasmid expressing myc-tagged N from SARS-CoV-2 delta variant, delta variant-specific mutation sites D63G, R203M and D377Y were included in the primers to amplify three fragments with overlapped ends from pCW57.1-SARS-CoV-2 N (Wuhan strain). These three fragments were seamlessly ligated in one step through NEBuilder HiFi DNA Assembly Kit according to manufacturer’s instructions. To generate lentivirus plasmid expressing myc-tagged N from SARS-CoV-2 omicron variant, omicron-specific mutation sites P13L, R203K and G204R were introduced into pDONR221-N through NEBuilder HiFi DNA Assembly to make an intermediate mutant construct, pDONR221-N mutant. The del31-33 was introduced into pDONR221-N mutant which has P13L, R203K and G204R already, through amplification with a pair of primers missing Glu-Arg-Ser on each 5’ terminal end by Q5® Site-Directed Mutagenesis Kit. The omicron variant N was finally cloned into Gateway-compatible pCW57.1 or pcDNA3.1 vectors by using Gateway™ LR Clonase™ II Enzyme mix.

To construct lentivirus plasmid expressing myc-tagged N of SARS-CoV-1 or MERS-CoV, the gateway entry plasmids, pEntry-SARS-CoV-1-N and pEntry-MERS-CoV-N were obtained from Addgene (#168852 and #168833). The Kozak sequence and myc tag were inserted into the N terminal ends of the N proteins through integration into the primers to amplify two fragments with overlapped ends for NEBuilder HiFi DNA Assembly. The myc-tagged SARS-CoV-1 and MERS-CoV N proteins were later cloned into pCW57.1 through Gateway™ LR Clonase™ II Enzyme mix.

To construct plasmid expressing cGAS tagged with 3xFlag, the cGAS was cloned from the total RNA from THP-1 cells and inserted into pcDNA3.1-3xFlag vector through NEBuilder HiFi DNA Assembly Kit. To construct plasmid expressing PABP1 tagged with 3xFlag, the PABP1 was moved from an existing pcDNA3.1-myc vector based construct to pcDNA3.1-3xFlag vector through restriction sites, NotI and XbaI. The pcDNA3.1 plasmid expressing V5-tagged IGF2BP3 was previously constructed by Dr. Serina Huang in our lab.

### Generation of stable HEK293T inducible SARS-CoV-2 protein expression cell lines

To generate inducible HEK293T cell lines, we utilized the enhanced PiggyBac (ePB) transposable element system, generously provided by the Brivanlou laboratory at Rockefeller University, as previously described^62^. Parental wildtype HEK293T cells were co-transfected with an equal ratio of the transposase plasmid and the ePB transposon vector encoding one of five SARS-CoV-2 proteins; N, membrane (M), non-structural protein (nsp) 14, nsp15, and nsp16, previously identified to be innate immune antagonists. Forty-eight hours post-transfection, cells were subjected to selection with 1.5 μg/mL puromycin to enrich for a population of HEK293T cells carrying stable, inducible expression of N, M, nsp14, nsp15, or nsp16.

### Cloning of an NF-κB responsive luciferase reporter plasmid

Previous work established the requirement of 5 tandem NF-κB binding sites before a promoter for the most efficient reporter activity when tracking NF-κB^63^. Additionally, the NF-κB consensus binding site was determined to be GGRNNNNYCC^64^. Therefore, we constructed a gene block containing 5 tandem NF-κB binding sites and the −56 to +20 region of an NF-κB regulated gene, flanked by EcoRI (5’) and NheI (3’) cut sites along with four random additional nucleotides on either end (5’-AGGCGAATTCGGAATTTCCCGGGAATTCCCGGGATTACCCGGGATTTTCCGGGATTTCCCTCTGAATAGAGAGAGGACCATCTCATATAAATAGGCCATACCCATGGAGAAAGGACATTCTAACTGCAACCTTTCGCTAGCGGCC-3’). This gene block was initially cloned into the pCR-Blunt II-TOPO vector using the TOPO Blunt cloning kit (ThermoFisher) for amplification. Both the TOPO vector containing the gene block and the pGL3-IFNB1-FLuc plasmid (AddGene #102597) were digested using EcoRI and NheI (New England Biolabs). The appropriate fragments were isolated from agarose gels and ligated together using the T4 ligation reaction kit from New England Biolabs. Chemically competent bacteria were then transformed with the ligation mixture followed by antibiotic selection.

### IFN and NF-kB activation luciferase assays

Inducible HEK293T cells were seeded at a density of 1×10^5^ cells/mL in DMEM supplemented with 10% FBS and 1µg/mL of doxycycline. The following day, the cells were transfected using the TransIT-mRNA transfection kit (MirusBio) with a mix of 100 ng of plasmid expressing firefly luciferase under control of the IFNB1 promoter or plasmid expressing firefly luciferase under control of NF-kB binding sites, 10 ng of plasmid constitutively expressing renilla luciferase under control of the CMV1 promoter, and 500 ng of poly I:C (for the IFNB1 reporter) or 20ng/mL of recombinant TNF-a (for the NF-κB reporter). Twenty-four hours after transfection, cells were harvested by aspirating the media and dispensing 100µL of 1X passive lysis buffer (Promega). The dual luciferase kit was used to determine firefly and renilla luciferase activities according to manufacturer’s instructions (Promega). Cell lysates supplemented with luciferase substrates were analyzed for luciferase activity using a BioTek plate reader.

### Lentivirus packaging and stable THP-1 cell line construction

To package lentivirus, Lenti-X HEK293T cells (Takara) were transfected with pMD2.G (Addgene, # 12259), psPAX2 (Addgene, #12260) and pCW57.1-N plasmid of different coronaviruses. The supernatant samples containing lentivirus particles were collected 48 hours post transfection and concentrated through Lenti-X™ Concentrator according to manufacturer’s instruction.

Spinoculation was performed for lentivirus transduction in THP-1 cells to stably integrate the N protein in the genome. Briefly, 3 X 10^5^ THP-1 cells/well were seeded in 500 ml transduction medium (RPMI 1640 with 5% FBS and 4 μg/ml polybrene) with 100 ml concentrated lentivirus in a 12-well plate. The THP-1 cells were spinoculated at 1500 rpm at 37°C for 1 hour and recovered in incubator for 4 hours before 500 μl of RPMI 1640 complete medium was added to each well. The transduced THP-1 cells were expanded for 48 hours and added with 2 μg/ml puromycin for 5-day selection. The stable THP-1 cell lines were maintained in RPMI 1640 with 10% FBS, 1X NEAA, 1X P/S and 0.2 μg/ml puromycin.

### Macrophage stimulation with TLR and RLR agonists and supernatant cytokine and chemokine detection

For PRR stimulation, the macrophages derived from stable THP-1 cell lines or parental THP-1 cells were treated with 2 μg/ml doxycycline in RPMI 1640 supplemented with 10% human AB serum, 1X NEAA, 1X P/S for 48 hours to induce N expression or serve as controls. The doxycycline-treated macrophages were then stimulated with 1 μg/ml CL097, an imidazoquinoline compound, to stimulate the TLR7/8 receptor, or 10 μg/ml poly (I:C) to stimulate the TLR3 receptor in RPMI 1640 supplemented with 5% human AB serum and 2 μg/ml doxycycline. For RLR stimulation, the final concentration of 2 μg/ml poly (I:C) was transfected into doxycycline-treated macrophages through lipofectamine RNAiMAX (Invitrogen) according to manufacturer’s instructions. Four hours after PRR stimulation, the cells were harvested for downstream processing.

For supernatant cytokine and chemokine detection, the culture mediums were collected after PRR stimulation, followed by a centrifugation at 200 g at 4 degree for 5 min to remove the cell debris. The clarified supernatants were then submitted to Eve Technologies (Canada) for pro-inflammatory/inflammatory cytokine and chemokine detection.

### RNA isolation for RNA-seq analysis

The parental and N-expressing THP-1 cells were seeded in biological duplicates at the density of 5 × 10^5^ cells/well. After differentiation, doxycycline treatment and PRRs stimulation, the cells were lysed in TRizol reagent (invitrogen), followed by total RNA extraction with Direct-zol™ (Zymo). The purified RNAs were submitted to BGI for transcriptome sequencing.

### RNA-seq bioinformatic analysis

The RNA-seq data were first processed to remove adapter sequences and low-quality reads using Trimmomatic, followed by the pseudoalignment to the human genome GCHr38 using Salmon (v1.10.2)^65^. The read counts of genes were generated by Tximeta^66^, and the differential expression tests between the groups of interest were analyzed by edgeR^67^ under the GLM framework. The similarities of the samples were explored by generating multi-dimensional scaling (MDS) plots through edgeR, and the visualization of MDS plot was further polished through ggplot2. For gene ontology (GO) and pathway analysis, the selected differentially expressed genes (DEGs) were submitted to DAVID database^68^ or Enrichr^38^ to search for over-represented GO and pathway terms. For transcription factor prediction, the selected DEGs were submitted to the Erichr to search most relevant regulatory transcription factors in TRRUST^37^ database. The volcano plots of gene expressions in N-expressing macrophages, the histograms of delta variant fold changes of N DEGs, the bar charts of the GO analysis results and the lollipop plots of transcription factor predictions were all visualized by corresponding functions in ggplot2^69^. The Venn diagrams were generated through ggVennDiagram^70^, and the giant heatmap of overlapped DEGs in all treatment conditions were plotted through pheatmap^71^.

### Reverse transcription and qPCR

To analyze intracellular cytokine and chemokine expressions after SARS-CoV-2 infection, VLP inoculation and PRR stimulations, the total cellular RNAs were isolated as previously described and reverse transcribed into cDNAs through the Protoscript II First Strand cDNA Synthesis Kit (NEB) according to manufacturer’s instruction. The interested cytokine and chemokines were detected with NEB Luna qPCR dye with specific primers from PrimerBank^72^. The qPCR reactions were run on CFX Opus system (Rio-Rad) with the cycling conditions suggested by NEB. The relative fold changes of the cytokines and chemokines of infected or stimulated samples to untreated controls were calculated by 2 (-Delta Delta C(T)) method.

### Immunoprecipitation and immunoblotting

To prepare for co-immunoprecipitation for affinity purification-mass spectroscopy(AP-MS), 6 x 15-cm dishes of parental and myc-tagged SARS-CoV-2 N-expressing (delta or omicron variant) THP-1 cells were differentiated into macrophages with 2 × 10^7^ cells per dish. Half of the cells per dish were treated with 2 μg/ml doxycycline to induce N expression or serve as drug-treated controls. The other half of the cells were maintained in RPMI 1640 with 10% human AB serum, 1X P/S without doxycycline treatment. The cell culture scale for AP-MS validation is smaller, the THP-1 culture and differentiation for immunoprecipitation were carried out in 6-well plates with 1.2 × 10^6^ cells seeded in each well. Forty-eight hours later, cells in each dish were lysed with 1% NP40 lysis buffer (ThermoFisher Scientific) supplemented with 1X Phosphatase Inhibitor Cocktail I (abcam), 1 μM DDT, and Complete EDTA-free protease inhibitor mixture tablet (Roche). Cell lysate from each dish was centrifuged at 14000 rcf for 15 mins. The clarified supernatants were incubated with anti-myc agarose beads (EZview™ Red Anti-c-Myc Affinity Gel, Millipore) for 45 min at 4°C. After washing with NP40 lysis buffer 4 times, proteins were eluted with urea buffer (8M urea, 100 mM Tris HCl (pH 8)) for mass spectrometry analysis or immunoblot validation.

For immunoblot, proteins were resolved by SDS-PAGE in 4-15% precast Mini-PROTEAN TGX Gels (Bio-Rad) in conventional Tris/Glycine/SDS buffer and later blotted to PVDF membrane (Bio-Rad), followed by the detection with primary antibodies and HRP-conjugated secondary antibodies. Immunoblots were imaged by chemiluminescence with the ProSignal Pico ECL Reagents (Genesee Scientific) on a ChemiDoc (Bio-Rad). The antibody information is in **Supplementary table 5**.

### AP-MS and proteomic data visualization

Following the affinity purification, samples were reduced with tris-(2-carboxyethyl) (TCEP) (10 mM final) for 30 min at RT and alkylated with 2-chloroacetamide (40 mM final) at RT for 30 min, sequentially, with shaking on a ThermoMixer (Eppendorf) at 1200 rpm. Next, Tris-HCl buffer (0.1 M, pH 8) was added to the samples to dilute the urea to 1 M final concentration, and 2 μg of sequencing grade Trypsin (Promega) and 1 μg of LysC (Wako) was added to the samples and digested overnight on a ThermoMixer at 1200 rpm at RT. Peptides were acidified with trifluoroacetic acid (0.5% final concentration) and desalted with spin desalting columns (Higgins Analytical) and eluted with 50% acetonitrile (ACN), 0.1% formic acid. The eluted peptides were dried in a SpeedVac (Labconco) and resuspended in 50 μL of 0.1% formic acid. Dried peptides were resuspended in 0.1% FA in LC/MS grade and analyzed on a timsTOF HT mass spectrometer (Bruker Daltonics) paired with a Vanish Neo UHPLC system (ThermoScientific). Mobile phase A consisted of 0.1% FA in MS grade water, and mobile phase B consisted of 0.1% FA in 100% MS grade ACN. Liquid chromatography was performed in a trap-and-elute mode. First, peptides were trapped on a PepMap Neo trap column (5 mm, 100 Å pore size, 5 μm particle size). Then, they were separated by reversed-phase chromatography on an Aurora Elite C18 reverse phase column (15 cm length, 75 μm diameter, 1.7 μm particle size for captive spray, IonOptiks). The 45-minute LC gradient was run with a flow rate of 300 nL/min set as follows: 5-35% B over 37 min, then to 45% B over 4 min, then to 60% B over 1 min, and then to 95% B for 3 minutes. The column was maintained at 50°C using a column oven for Bruker Captive Spray source (Sonation Lab Solutions). Ionization was performed using a CaptiveSpray source (Bruker Daltonics) at 1700 V. On the timsTOF HT, equal-size windows of 25 Da were designed with an overlap of 1 Da to maximize the precursors ion coverage for further MS/MS. The ion accumulation time and ramp times in the dual TIMS analyzer were set to 100 ms each. In the ion mobility (1/K0) range 0.6 to 1.6 Vs cm-2, the collision energy was linearly decreased from 59 eV at 1/K0 = 1.46 Vs cm-2 to 20 eV at 1/K0 = 0.62 Vs cm-2 to collect the MS/MS spectra in the mass range 265.0 to 1370.0 Da. The estimated mean cycle time was 1.59 s.

The raw files were processed with Spectronaut (Biognosys, ver. 19.0) with the directDIA+ (Deep) search algorithm. Carbamidomethylation (cysteine) was set as a fixed modification for database search. Acetylation (protein N-term), oxidation (methionine), and phosphorylation (serine, threonine, tyrosine) were set as variable modifications. Reviewed human proteome and SARS-CoV-2 protein sequences (downloaded from UniProt) were used for spectral matching. The false discovery rates for the PSM, peptide, and protein groups were set to 0.01, and the minimum localization threshold for PTM was set to zero. For MS2-level area-based quantification, the cross-run normalization option was unchecked (normalization was performed later using MSstats), and the probability cutoff was set to zero for the PTM localization. Quantitative analysis was performed in the R statistical programming language (v.4.4.1). Initial quality control analyses, including inter-run clustering, correlations, principal component analysis (PCA), peptide and protein counts, and intensities were completed using custom R code. Statistical analysis of phosphorylation and protein abundance changes between exposed and control samples were computed in MSstats (ver 4.16.0)^73^. MSstats was parameterized to perform normalization by median equalization, no imputation of missing values, and median smoothing (Tukey’s median polish) to combine intensities for multiple peptide ions or fragments into a single intensity for their protein group, and statistical tests of differences in intensity between conditions. Default settings for MSstats were used for adjusted P values. By default, MSstats uses the Student’s t-test for P value calculation and the Benjamini– Hochberg method of FDR estimation to adjust P values. Identified proteins were subjected to protein– protein interaction scoring using SAINTexpress (http://apostl.moffitt.org/)^74^. Protein interactions with a Bayesian false-discovery rate (BFDR) ≤0.05 and average spectral count ≥11 were selected as high-confidence protein-protein interactions and visualized with Cytoscape (ver 3.10.3)^75^.

### Measurement of Transendothelial electrical resistance (TEER)

TEER measurements of hPSC-BMELCs or HCAECs in a transwell system were conducted using the EVOM3 (World Precision Instruments). STX2 Plus electrodes were positioned with one prong in the apical chamber and the other in the basolateral chamber, ensuring both were submerged in media at an approximate depth of 0.5 cm in a 12 well plate. Readings were taken starting once daily after cell seeding, with resistance values recorded alongside a cell-free control well containing plain media, which was used for background subtraction. Cell media was changed daily, and for treatment with macrophage supernatant, media was replaced with the cells’ respective medium with 20% macrophage supernatant. TEER in Ωcm² was calculated by taking the corrected resistance values (Ω) and multiplying by the surface area (cm²) of the transwell insert, providing a measure of barrier integrity.

### Statistical analysis

Bar graphs with appropriate statistical analyses were generated and performed by Graphpad Prism 10. Representative results from at least 2 independent experiments are shown as means ± standard deviation (SD). Statistical significance was indicated in figures as: ∗ (p < 0.05), ∗∗ (p < 0.01), ∗∗∗ (p < 0.001) and ∗∗∗∗ (p ≤ 0.0001).

---

**Supplementary Figure 1.**
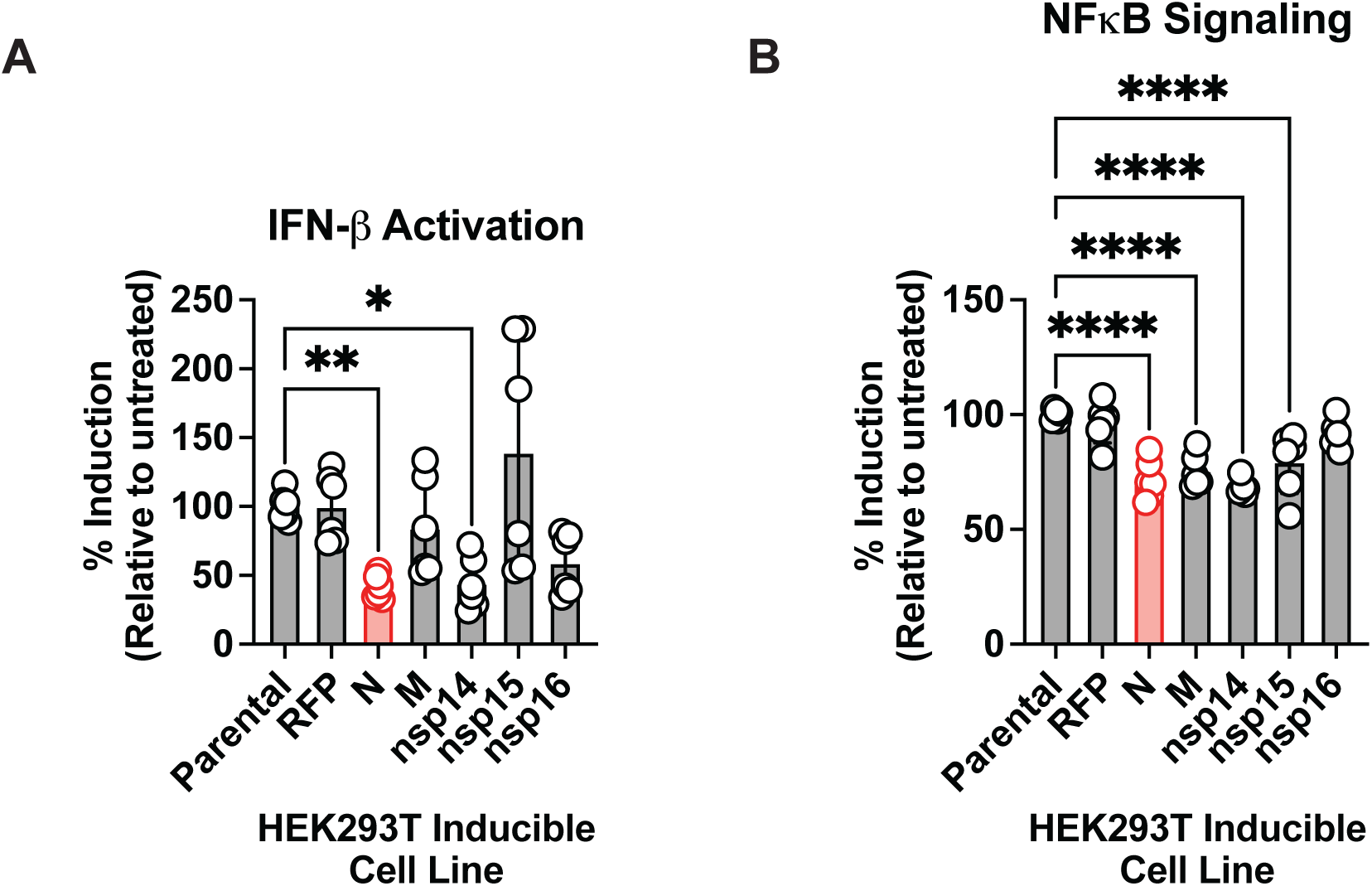
**(A-B).** The inducible HEK293T cell lines were treated with 1 mg/ml of doxycycline (dox) to express different SARS-CoV-2 proteins from Wuhan strain (N, M, nsp14, nsp15, nsp16). The parental cells and a dox-inducible HEK293T cell line expressing RFP were served as controls. After 24-hour induction with dox, the cells were transfected with a IFNB1 reporter plasmid (A) or NF-kB reporter plasmid (B) which expresses firefly luciferase either under the control of IFNB1 promoter or NF-kB binding sites. At the meantime, the cells were co-transfected with a renilla luciferase plasmid for normalization. Cells were either transfected with poly(I:C) (A) or treated with TNF (B) at the same time as plasmid transfection. Asterisks indicate statistically significant differences in IFN induction or NF-kB signaling relative to parental cells (two-way ANOVA and Šidák’s multiple comparisons): *, p <0.05; **, p < 0.01; ****, p < 0.0001. Non-significant differences are not shown.

**Supplementary Figure 2.**
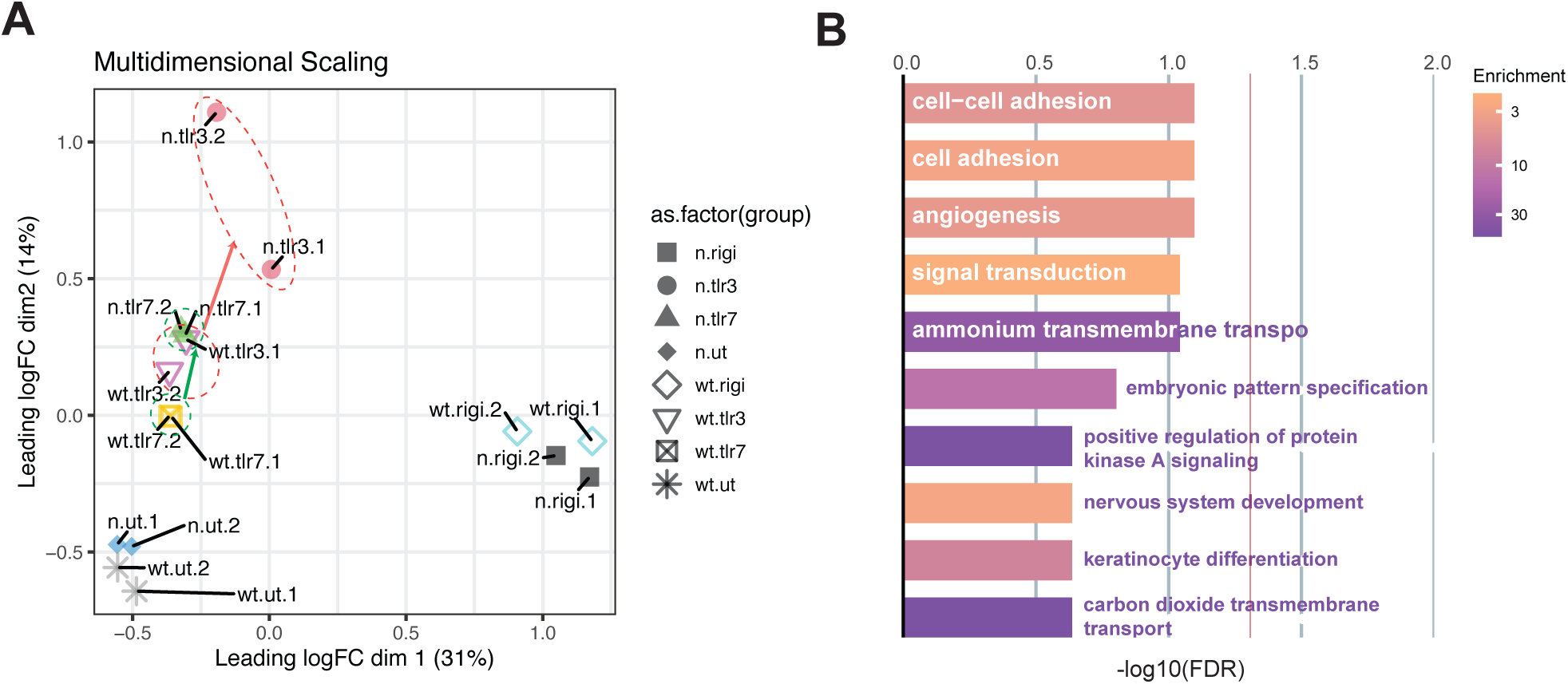
**(A).** The multidimensional scaling analysis (MDS) plot of RNA-seq samples in biological duplicates. Each dot represents one RNA-seq sample. The distances between the dots summarize the gene expression differences between the samples. **(B).** The biological process (BP) analysis of N-upregulated DEGs in RIG-I/MDA5 stimulations. Top 10 enriched gene ontology terms of 300 N-upregulated DEGs (delLog2FC>1) in macrophages after RIG-I/MDA5 stimulation. None of the bars exceed -log10(FDR) cutoff (red line, FDR=0.05).

**Supplementary Figure 3.**
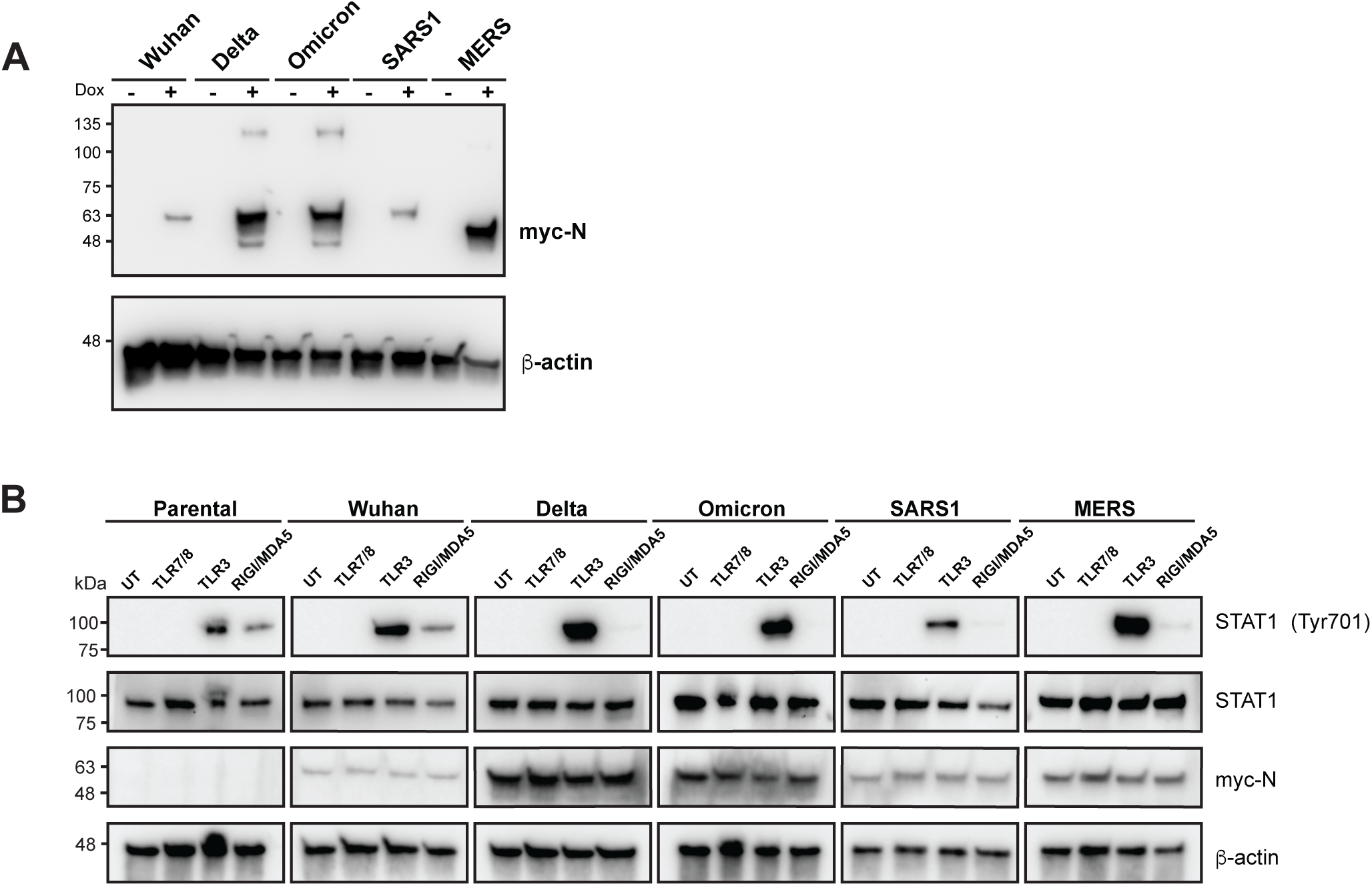
**(A).** Western blot validation of dox-induced N expressions from different THP-1-derived macrophage cell lines. The THP-1 derived macrophage which can inducibly express the N protein of SARS-CoV2 Wuhan strain, SARS-CoV2 delta variant, SARS-CoV2 omicron variant, SARS-CoV1 or MERS was left untreated or stimulated with dox (2 mg/ml) for 48 hours. All the N expressions were detected and evaluated through its N-terminally-fused myc tag. **(B).** The phosphorylation of STAT1 influenced by different coronavirus N proteins after PRR stimulations. As previously mentioned, the THP-1 derived stable macrophage cell lines and parental macrophages were treated with dox (2 μg/ml) for 48 hours to express the N proteins from different SARS-CoV2 variants and coronaviruses or serve as a control (parental). The cells were sequentially stimulated with different PRR agonists for 4 hours to activate TLR7/8, TLR3 and RIG-I/MDA5. The cell lysates were then collected to detect the phosphorylation at Tyr701 in STAT1 by western blot.

## Supplementary Tables

**Table 1: N DEGs for bar graphs in Fig. 2D-2F**

**Table 2: Overlapped DEGs for heatmap in Fig. 2G**

**Table 3: Host factor functional annotation for Fig. 4C**

**Table 4: Macrophage cytokine secretion for Fig. 5A-5E**

**Table 5: Primers and antibodies**

